# Growth rate and yield are not constrained by a trade-off in microbes

**DOI:** 10.64898/2026.09.08.750265

**Authors:** Ziyin Jasmine Xiong, Jana S. Huisman, Jeff Gore

## Abstract

Ecological theory traditionally posits widespread trade-offs in organismal growth, survival, and reproduction. One core trade-off is the relation between maximum intrinsic growth rate (*r*) and carrying capacity (*K*), proposed as a key mechanism explaining species coexistence and the maintenance of biodiversity. To quantify how *r* relates to *K* in microbes, we analyzed more than 70,000 microbial growth curves in dozens of media conditions. In this set, we observed a strong positive correlation between *r* and *K*, challenging classic life-history expectations of a trade-off. This unexpected “trade-up” can be recovered by a model assuming a non-zero maintenance cost in cells. The model predicts a stronger trade-up under stress conditions that reduce growth rate, consistent with empirical data showing that salinity stress strengthens the trade-up. We also found that community assembly weakens but does not eliminate this positive *r*-*K* relationship. The robust growth-yield trade-up between species contrasts with patterns within species. Single-gene knockout datasets in *Escherichia coli* and budding yeast show no or only weak *r*–*K* relationships. Together, our results suggest that the lack of strong trade-offs between growth rate and carrying capacity may be widespread across species, emphasizing the importance of alternative trade-offs and coexistence mechanisms beyond the classical *r*-*K* framework.

## Introduction

Ecological theory has long emphasized trade-offs as a central mechanism promoting coexistence [1, 2, 3]. Trade-offs arise when improvement in one trait leads to a deterioration in another trait, for example, mobility and energy conservation during animal migration, or offspring survival and number [4, 5, 6, 7]. A particularly fundamental trade-off is the proposed relationship between maximum intrinsic growth rate, *r*, and carrying capacity, *K*. This trade-off has been widely invoked in life-history theory and community ecology because trade-offs among metabolic or demographic traits prevent a single optimal species from dominating a community and thereby promote biodiversity [1, 8, 9, 10, 11, 12]. Yet whether growth rate and carrying capacity generally trade off across species remains unresolved.

Microbes provide a powerful system for testing the existence of a trade-off between *r* and *K* because both traits can be measured quantitatively across many taxa and environments [13]. In microbial batch growth with a fixed initial resource supply, *K* corresponds to biomass yield. We therefore refer to this relationship interchangeably as the *r*–*K* or growth rate–yield relationship. Microbial communities in snow-covered soil exhibit evidence of a rate-yield trade-off [14], as do chemostat experiments of microbial communities and studies of yeast metabolism within strains [15, 16, 17]. This is consistent with thermodynamic and metabolic arguments which also predict a rate-yield trade-off, particularly between fast but inefficient fermentation and slower but higher-yield respiration [18, 19]. However, these datasets are often limited in size, condition-specific, or based on indirect comparisons among nutrient consumption, respiration, and growth efficiency [17, 20, 21, 22]. Indeed, studies of mixed aquatic bacterial communities provide contrasting evidence: bacterioplankton growth rate and growth efficiency can increase together as nutrient availability rises [23, 24, 25]. Yet, these studies examine community-level responses rather than comparisons among individual taxa. More recently, a study of 23 soil microbial strains found no relationship between maximum growth rate and growth efficiency, with efficiency instead varying primarily with resource and species identity [26]. The broad pattern of *r* and *K* across microbial species therefore remains uncertain [14, 27].

The expectation of an *r*–*K* trade-off is largely a legacy of life-history theory rather than a requirement of the models that define the two traits. In the logistic model, carrying capacity is simply the inverse of the strength of intraspecific density dependence [27]. Classical *r*- and *K*-selection theory contrasts selection at low density, which favors high intrinsic growth rate, with selection near equilibrium, which favors performance under strong density dependence [2], although competitive ability at high density is not generally captured by monoculture *K* alone [28]. The framework does not inherently require a negative relationship between *r* and *K*, but it has nevertheless given rise to the expectation that fast-growing species reach lower carrying capacities, and vice versa [29]. Mechanistic models that derive both traits from underlying physiology provide one route to addressing this question, although the sign of the *r*–*K* relationship has so far remained unresolved. Existing models link growth rate and yield through detailed cellular constraints or evolutionary dynamics [30, 31, 32], while maintenance-energy models show that non-growth-associated costs affect both traits without specifying their relationship [33, 34, 35]. This motivates us to combine modeling with experimental data to determine the relationship between *r* and *K*.

Here, we determine the relationship between growth rate *r* and carrying capacity *K* across natural microbial species. We analyze over 70,000 growth curves from phylogenetically diverse aquatic bacteria grown under many environmental conditions to uncover broad patterns of *r* and *K*. Contrary to expectations, we find a robust and positive relationship, a “trade-up”, between growth rate and carrying capacity. To explain this pattern, we develop a minimal resource-uptake model with maintenance costs in which both *r* and *K* emerge from the growth dynamics rather than being imposed as model parameters. The model predicts conditions that strengthen the trade-up, and we experimentally demonstrate that stress increases its strength while community assembly weakens it. Collectively, our results show that the lack of a trade-off between growth rate and carrying capacity may be widespread across natural microbial taxa and microbial model organisms.

## Results

### A positive *r* − *K* relationship persists across diverse environments and taxonomical levels

In this study, we set out to test the relationship between maximum intrinsic growth rate (*r*) and carrying capacity (*K*) in natural microbial species. For our primary dataset, we used extensive growth measurements collected in the context of our recent study on the impact of environmental (salinity) stress on aquatic microbes [36]. In total, the dataset contains growth measurements for 103 isolates from 55 species, isolated from three environments (freshwater, estuarine, and marine habitats; Fig. 1A). These isolates represent a phylogenetically diverse group, representing 5 classes, 11 orders, 14 families, and 29 genera. Each isolate was grown in rich medium (marine broth) at 12 salinities for 48 hours with growth monitored by optical density at 600 nm (OD_600_) as an established proxy for population density [37]. The resulting growth curves were used to estimate the maximum intrinsic growth rate (*r*) and carrying capacity (*K*). We considered whether *r* and *K* show a “trade-up” (positive relationship), no relationship, or a trade-off (Fig. 1B). Across this diverse set of aquatic microbes, we observed a strongly positive relationship between *r* and *K*, indicating a pervasive trade-up (Fig. 1C). This trade-up persists when analyzing species at the salinity associated with their isolation habitat (freshwater, estuary, or marine; Fig. 1D) and all other salinities (Extended Data Fig. 1). To avoid an artificial positive relationship driven by data points at (0, 0) (i.e., *r* = 0 and *K* = 0), we excluded isolates with no detectable growth, as points that are simultaneously low in both traits can inflate the apparent positive association between *r* and *K*. Our observation of a robust growth-yield trade-up is surprising because it breaks the common expectation of a trade-off, in which faster-growing species are expected to be less efficient in resource utilization and reach lower carrying capacities.

**Figure 1.**
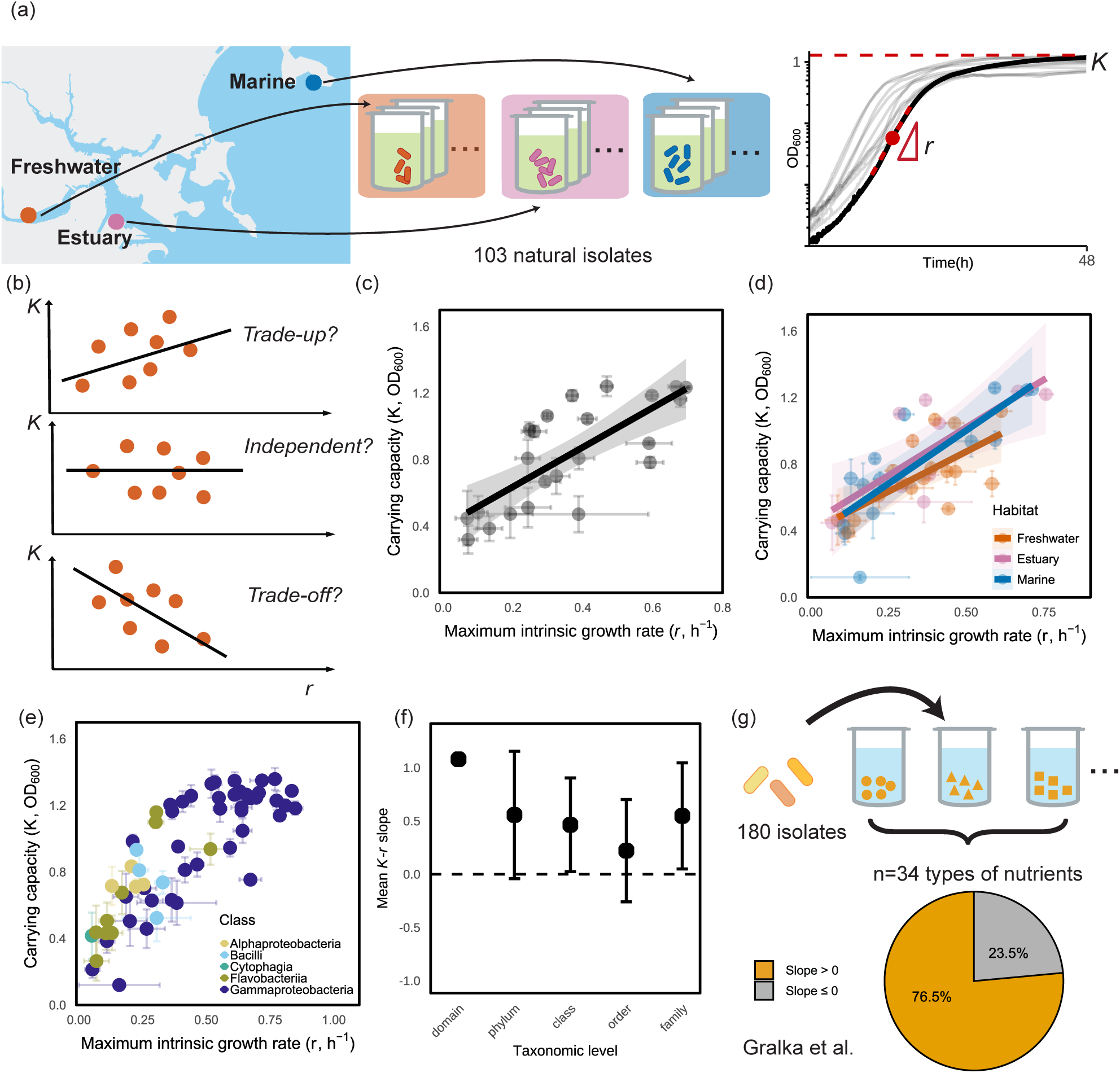
A positive *r*–*K* relationship is widespread among microbes sampled from diverse ecological habitats. **(a)** Schematic of the isolate sampling and growth rate measurement protocol of our primary dataset [36]. Aquatic microbial communities were collected from three sites around Boston, MA, representing freshwater, estuary, and marine habitats. Population growth curves were used to estimate the maximum intrinsic growth rate *r* and carrying capacity *K* (maximum optical density). **(b)** Possible relationships between maximum intrinsic growth rate *r* and carrying capacity *K*: a positive trade-up, no relationship, or a negative trade-off. **(c)** Growth rate and carrying capacity of aquatic microbial species grown at a salinity of 30 g L*^−^*^1^. Each point represents a species (*n* = 23), with error bars showing the SEM of *r* and *K* across isolates within that species. The estimated *K*–*r* slope is 1.30 (SE = 0.24). **(d)** Growth rate and carrying capacity of aquatic microbial species grown at the salinity of their sampling location: Freshwater at 5 g L*^−^*^1^ (*n* = 17, slope 0.96 ± 0.27), Estuary at 30 g L*^−^*^1^ (*n* = 10, slope 1.15 ± 0.33), and Marine species at 35 g L*^−^*^1^ (*n* = 14, slope 1.29 ± 0.25). Each point represents one species, with isolates first grouped by habitat and then averaged by species within each habitat. Error bars show the SEM of *r* and *K* across isolates within that species. **(e)** Growth rate and carrying capacity of aquatic microbial isolates grown at 30 g L*^−^*^1^, colored by their taxonomic class. Each point represents one isolate, and error bars show the SEM of *r* and *K* across replicate measurements of that isolate. **(f)** Mean within-group *K*–*r* slope across taxonomic levels. At each taxonomic level, we calculated slopes within each group at that level (e.g. within all Flavobacteria). At least *n* ≥ 4 isolates were required to compute a within-group slope (total number of slopes: *n* = 1, 3, 4, 4, 5 for domain, phylum, class, order, and family respectively). Black points show the mean *K*-*r* slope ± SEM across groups at each taxonomic level. The dashed line indicates a slope of zero. **(g)** Analysis of the Gralka *et al.* [38] dataset, which measured growth of 180 microbial strains in different single-resource environments. We considered only resources on which at least 50 isolates grew (*n* = 34 resources).

Because evolutionary theory often emphasizes trade-offs among closely related organisms, we next asked whether the observed trade-up reflects broad differences between distant taxa or also persists within more closely related groups. We first assessed the *r*–*K* relationship within a taxonomic group. We grouped isolates by their taxonomic classification (at the family, order, class, or phylum level) and calculated the *K*–*r* slope within each group separately (obtaining e.g. 5 slopes for the families with *n >* 3 isolates in the dataset). The within-group mean *K*–*r* slopes were all positive, but tended to decrease toward lower taxonomic ranks (Fig. 1E,F). Sensitivity analyses showed that variation in the number of data points across groups did not impact this result (Supplementary Fig. 1). Second, we tested whether the trade-up persisted between data aggregated at different taxonomic levels. Specifically, we averaged the measured *r* and *K* values of isolates belonging to the same species, genus, family, order, or class, and then calculated the *r*–*K* relationship across these aggregated values. The positive *r*-*K* relationship remained evident (Extended Data Fig. 2). Together, these taxonomic analyses show that the positive *r*–*K* relationship is robust across taxonomic scales and is not driven by a particular subset of isolates.

We wondered whether these positive *r*-*K* relationships would extend beyond our primary dataset [36], and in particular whether they would arise during growth in different well-defined media. Because different resources impose distinct metabolic constraints on growth and biomass production, the relationship between *r* and *K* may depend on resource identity. We turned to another independent dataset which profiled 180 slow-growing marine bacterial strains across 34 individual carbon substrates [38]. Growth was monitored for 17 days, and *r* and *K* were estimated by fitting a growth curve for each strain–carbon source combination. We estimated the *r*–*K* relationship across isolates independently for each carbon source and quantified the corresponding slope. Although individual carbon-source-specific slopes were not generally statistically significant on their own, 76.5% of the resources showed positive slopes, and this fraction was significantly greater than expected by chance (two-sided exact binomial test, *P* = 0.0029). This indicates that positive *r*–*K* associations are common across carbon sources and are not specific to our primary salinity-gradient dataset (Fig. 1G). Together, these results show that aquatic microbial species consistently exhibit a positive *r*–*K* relationship, revealing a widespread trade-up.

### A minimal model incorporating cellular maintenance costs can explain the positive *r*–*K* relationship

To understand why our experimental results deviate from the classic expectation of a *r*-*K* trade-off, we developed a minimal model linking growth rate to carrying capacity. In our model, cells uptake resources from the environment, and the resulting intracellular resource pool is used for cell growth and division. In addition, there is a constant maintenance rate *M* that captures resources that are not used for growth but are instead dedicated to keeping the cell viable [33, 39, 40]. In the absence of a maintenance cost (*M* = 0), all resources are eventually converted into biomass and the final carrying capacity *K* is set by the total available resource and conversion efficiency (Extended Data Fig. 3D, E). A non-zero maintenance cost is sufficient to observe a trade-up between *r* and *K* (Fig. 2B, Extended Data Fig. 3).

**Figure 2.**
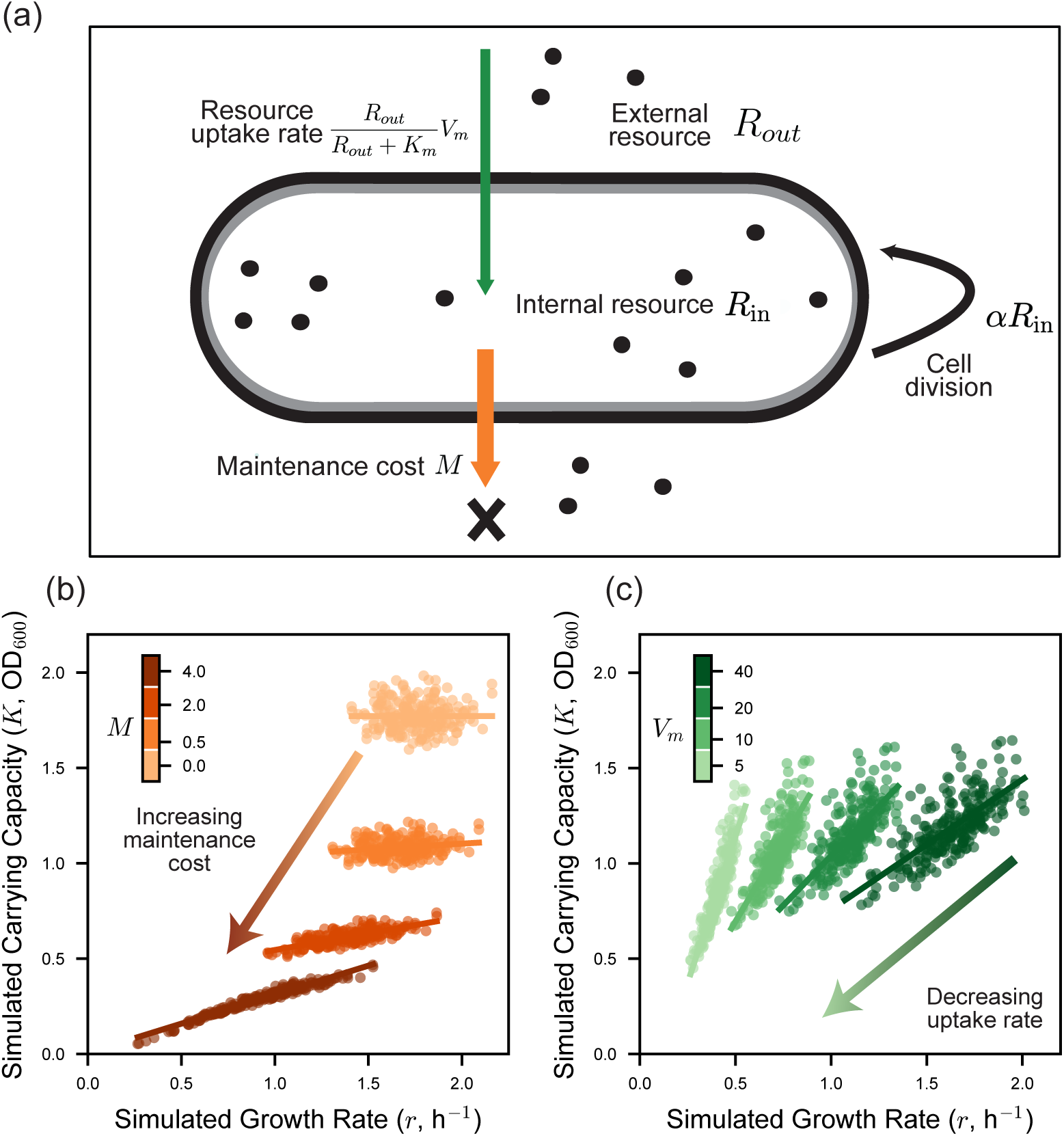
A minimal model incorporating cellular maintenance costs reproduces a positive *r*–*K* relationship. **a)** Schematic of the cell-level model including resource uptake, population growth, and maintenance costs. External resources *R_out_*are taken up into an intracellular resource pool *R_in_*via uptake rate 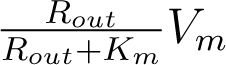. Internal resources are allocated to cellular maintenance *M* and population growth *αR_in_*. **b)** Simulation results from 1,000 runs in which the maintenance cost *M* was varied among 0, 0.5, 2.0, and 4.0 (light to dark orange). The parameters *C* and *V_m_* were sampled from normal distributions. A linear regression was fitted to estimate the *K*–*r* slope for each value of *M* . *M* = 0, slope = 0, *R*^2^ = 0; *M* = 0.5, slope = 0.06, *R*^2^ = 0.04; *M* = 2, slope = 0.17, *R*^2^ = 0.58; and *M* = 4, slope = 0.30, *R*^2^ = 0.96. **c)** Simulation results from 1,000 runs in which *V_m_* was varied among 5, 10, 20, and 40 (light to dark green). The parameters *α* and *M* were sampled from normal distributions. A linear regression was fitted to estimate the *K*–*r* slope for each value of *V_m_*. *V_m_* = 5, slope = 3.1, *R*^2^ = 0.83; *V_m_* = 10, slope = 1.7, *R*^2^ = 0.64; *V_m_* = 20, slope = 1.0, *R*^2^ = 0.57; and *V_m_* = 40, slope = 0.68, *R*^2^ = 0.55.

Specifically, as maintenance cost increases, the simulated relationship between *r* and *K* becomes progressively more positive (Fig. 2B). Mechanistically, the maintenance cost imposes a constant energetic demand that must be met before resources can be allocated to growth. Because this cost accumulates continuously over time, its effect depends on the speed at which a population depletes the external resource pool, creating a higher burden for slow than fast-growing bacteria. When *M >* 0, faster uptake of resources thus not only increases *r* but also *K*. These results show that a cellular maintenance cost alone is sufficient to generate the trade-up observed in the empirical data.

The model also predicts additional mechanisms that can strengthen the *r*–*K* trade-up. With maintenance costs present (*M >* 0), decreasing the maximum nutrient uptake rate *V_m_*strengthens the *r*–*K* trade-up (Fig. 2C, Extended Data Fig. 4A). This is because lowering *V_m_* slows resource uptake (lowering *r*) and prolongs the time required to reach stationary phase, which has a larger impact on *K* for species that continuously pay high maintenance costs (Extended Data Fig. 4B-E). Such reductions in resource uptake may arise naturally under environmental stress. For example, environmental stress can decrease cellular resource uptake efficiency through damage to transport machinery, altered membrane function, or diversion of energy away from nutrient uptake and toward stress response [41, 42, 43]. In particular, osmotic stress can rapidly inhibit active carbohydrate transport in *E. coli*, reducing carbon uptake even while cells remain viable and metabolically active [44]. Our model predicts that environmental conditions that reduce growth rates will strengthen the observed *r*–*K* trade-up.

### Increasing salinity stress strengthens the positive *r*−*K* relationship

To test the predictions of our model, we examined how increasing environmental stress affects the *r*–*K* relationship in experimental data. Our primary dataset includes growth measurements for each isolate across 12 salinity levels [36], allowing us to assess the impact of salinity on both *r* and *K*. For most species, both *r* and *K* are maximized near the salinity of the habitat from which the species were originally isolated [36]. Departures from this optimal salinity tend to decrease *r* and thus can be considered a physiological stress. Grouping species by their isolation habitat (freshwater, estuarine, and marine), we find that the slope of the *r*–*K* relationship across species was lowest near each habitat’s optimal salinity (Extended Data Fig. 5) and became progressively more positive as salinity increased beyond this optimum (Fig. 3A). Thus, increasing salinity stress strengthened the positive *r*–*K* trade-up.

**Figure 3.**
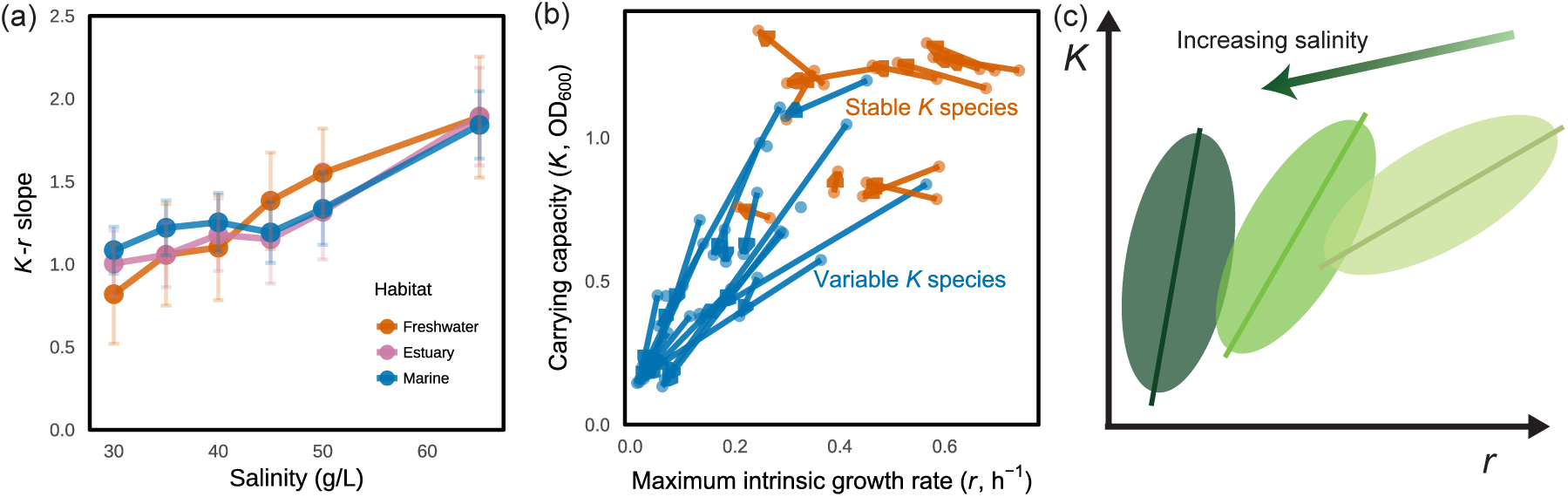
Increasing salinity stress amplifies the positive *r* − *K* relationship. **a)** Estimated slopes of carrying capacity *K* vs. maximum intrinsic growth rate *r* for species from three habitats measured across six salinities (30–65 g L*^−^*^1^). Points show the estimated *K*–*r* slope ± standard error. Colours indicate habitat: freshwater (orange; *n* = 19, 19, 18, 18, 18, 16 species), estuary (pink; *n* = 15, 15, 15, 15, 15, 13 species) and marine (blue; *n* = 25, 26, 27, 26, 26, 25 species). **b)** Orange arrows indicate trajectories in *r* − *K* space for species that maintain stable *K* (|Δ*K*| *<* 0.05 · *K_init_*) as salinity increases from 30 g/L to 65 g/L (*n* = 12). Blue arrows indicate trajectories for species with larger variation in *K* with increasing salinity (*n* = 22). **c)** Schematic illustration showing how species that experience reductions in *r* while maintaining stable *K* contribute to a steeper *r* − *K* relationship. In *r* − *K* space, species initially distributed in the lighter green region shift toward the darker green region as stress increases.

Consistent with the predictions of our model, this increase in *K*-*r* slope was primarily driven by species whose carrying capacities remained relatively stable with increasing salinity even as their growth rates decreased (Fig. 3B). To quantify this pattern, we classified species based on their change in carrying capacity Δ*K* = *K_end_* − *K_init_* under stress, defining as ‘stable’ those species with |Δ*K*| *<* 0.05 · *K_init_*. Mapping species trajectories in *r*–*K* trait space showed that these stable-*K* species predominantly started with higher *K* and *r*. This asymmetric response across species alters the distribution of *r* and *K* values. Growth rates become compressed while carrying capacities remain relatively broadly distributed. Consequently, the *r* − *K* distribution shifts toward a steeper positive relationship across species (Fig. 3C). As such, the empirical pattern mirrors the model prediction that stress strengthens the *r*–*K* trade-up.

### The *r*–*K* trade-up weakens after community assembly, consistent with generalized Lotka–Volterra (gLV) model predictions

We next asked how the *r*–*K* relationship changes when species interact within ecological communities. In natural environments, microbial species rarely grow in isolation, and interactions may select for species with specific *r*–*K* characteristics. Previous theoretical work based on generalized Lotka–Volterra (gLV) models predicts that extinction events during community assembly can increase the mean carrying capacity (*K*) and decrease the mean interaction coefficient (*α*) of the surviving species pool relative to the initial pool [45, 46]. Because these predictions concern *K* and *α* alone, it remains unclear what the relationship between *r* and *K* is for species growing in communities.

To examine how such ecological dynamics affect the *r*–*K* relationship, we constructed a gLV model with mortality and simulated community assembly (see Supplementary Methods and Extended Data Fig. 6). The mortality term represents dilution and other processes that remove individuals indiscriminately in laboratory or natural communities. In this framework, survival in an environment depends not only on interaction structure and carrying capacity, but also on whether species can grow fast enough to offset mortality. We found that, in the absence of mortality, community assembly does not generate a systematic relationship between *r* and *K* (Extended Data Fig. 6B-C). However, as mortality increases, assembly favors species with higher growth rates even when they have lower carrying capacities, and vice versa, leading to the emergence of a *r*–*K* trade-off, or a weakening of the trade-up if initially present within the species pool (Extended Data Fig. 6A,D,E). Community assembly under mortality therefore offers an alternative generator for *r*–*K* trade-offs.

To test these theoretical predictions, we made use of community propagation experiments from our primary dataset [36], in which natural communities were propagated in the laboratory for 2 weeks (Fig. 4A). Community composition over time was determined using 16S sequencing and matched to our isolate library, allowing us to link species identities and relative abundances to quantitative growth traits during community assembly (Supplementary Fig. 2). Consistent with predictions from the gLV model, we found that as species spent more time undergoing community assembly, the strength of the *r*–*K* relationship progressively decreased (Fig. 4B). Despite this reduction in slope, the *r*–*K* relationship remained positive throughout the assembly process (Fig. 4B, Extended Data Fig. 7). This pattern of a reduced *K*-*r* slope was robust across salinity treatments and across communities originating from different native habitats (Fig. 4C, Extended Data Fig. 8), indicating that the weakening of the relationship is a general consequence of community assembly rather than a feature of specific communities or environmental conditions.

**Figure 4.**
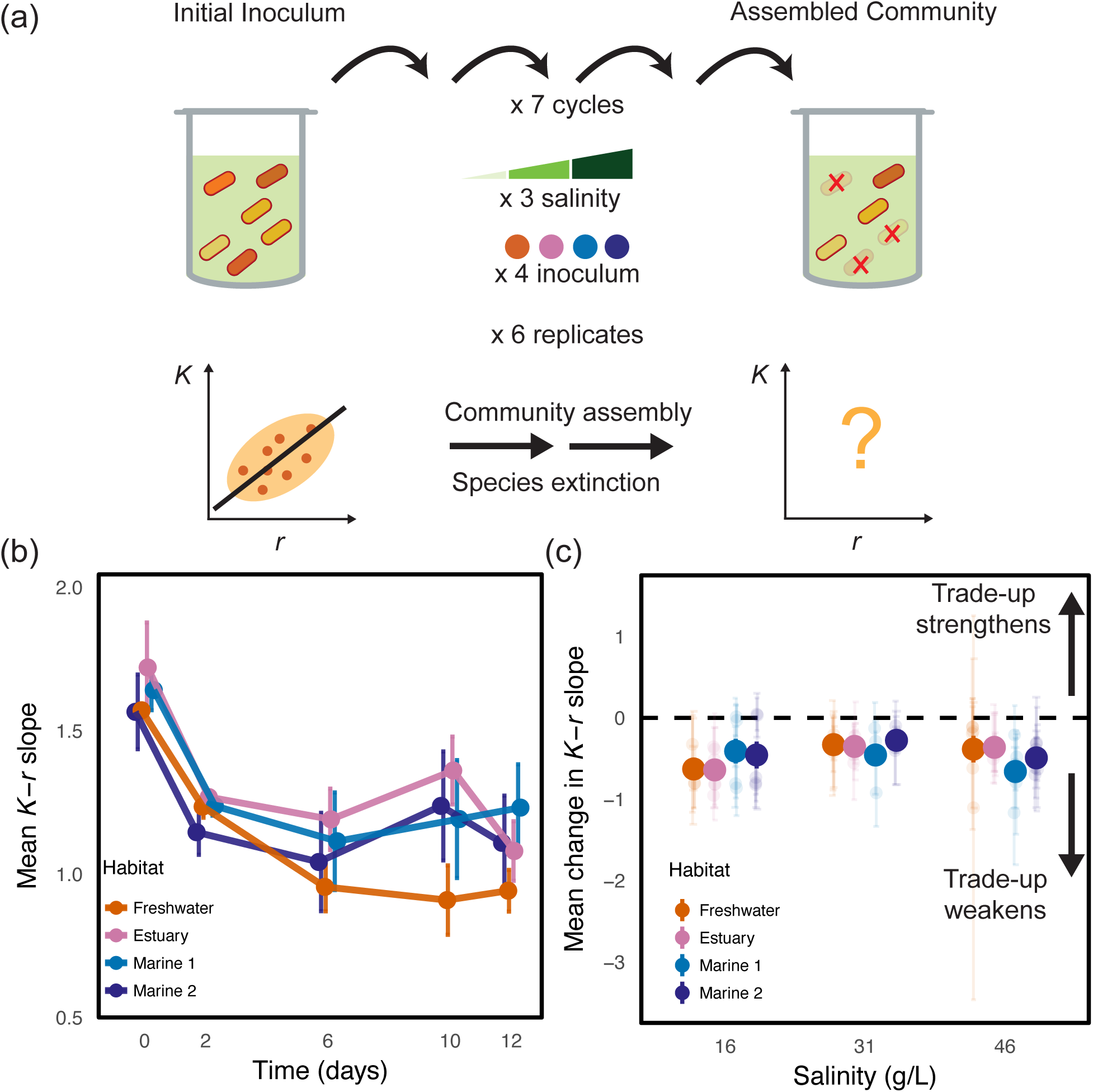
The *r*–*K* trade-up weakens after community assembly, consistent with generalized Lotka–Volterra (gLV) model predictions. **a)** Schematic of the experimental design: natural communities (4 inocula from different sampling locations) were propagated in the laboratory for 7 48-h cycles, at three salinity conditions (16, 31, 46 g/L). Each community and propagation condition has 6 replicates. **b)** Mean *K*–*r* slope ± S.E.M. for four natural communities measured over time during community assembly at 16 g/L salinity. The strength of the *r*–*K* relationship decreased over the course of community assembly. At Day 0, each point represents the mean slope of two communities per habitat. From Day 2 to Day 12, each point represents the mean slope of six communities per habitat. Most of the change in the species turnover occurred during the first two days after communities were brought into the laboratory, with modest changes thereafter. **c)** Mean change in the *K*–*r* slope between Day 0 and Day 12 of community assembly. The dashed line indicates 0 (no change). Across communities, the net change in slope is predominantly negative.

## Discussion

We set out to understand the relationship between *r* and *K* across microbial species, and found evidence of widespread trade-ups. Across two large datasets spanning diverse marine taxa, habitats, salinity conditions, and nutrient environments, we found that faster-growing species often also reached higher carrying capacities, in apparent contradiction to the classical expectation of a *r*–*K* trade-off. A minimal resource-uptake model suggests that this positive relationship between *r* and *K* can emerge from cellular maintenance costs: slow-growing populations lose more resources to maintenance than fast-growing populations. Consistent with this mechanism, increasing salinity stress strengthened the positive *r*–*K* relationship. Experimental community assembly weakened but did not eliminate this *r* − *K* trade-up. Together, these findings suggest that growth rate and carrying capacity are typically not constrained by a trade-off across species.

Although we observe a robust *r*–*K* trade-up across species, the same relationship is not necessarily expected during evolution within lineages. During evolution, genetic or physiological constraints may limit the combinations of *r* and *K* that a lineage can achieve. These constraints can create a boundary of possible trait combinations, or Pareto front, beyond which improving one trait requires sacrificing the other [47, 48, 49]. Different lineages may approach different Pareto fronts because of variations in their genetic backgrounds, metabolic pathways, and allocation of resources to growth and maintenance. A positive *r*–*K* relationship across species is therefore compatible with trade-offs within lineages, because ecological and evolutionary comparisons sample different sources of variation. Consistent with the possibility that *r*–*K* relationships differ across evolutionary scales, long-term evolution experiments in *E. coli* found negative *r*–*K* relationships among clones within 3 of 4 independently evolved populations, but no detectable relationship when comparing across populations [48]. In line with this distinction, we analyzed gene-knockout datasets in *E. coli* and budding yeast [50, 51, 52]. We find that genetic perturbations within the same strain display largely uncorrelated growth rate and carrying capacity (Fig. 5, Extended Data Fig. 9 - 10, Supplementary Fig. 3 - 4). Taken together, we conclude that there is typically a *r*–*K* trade-up between species but only weak *r*–*K* relationships within a species, highlighting the potential for scale-dependent relationships. However, in neither case do we observe evidence for widespread and strong growth-yield trade-offs.

**Figure 5.**
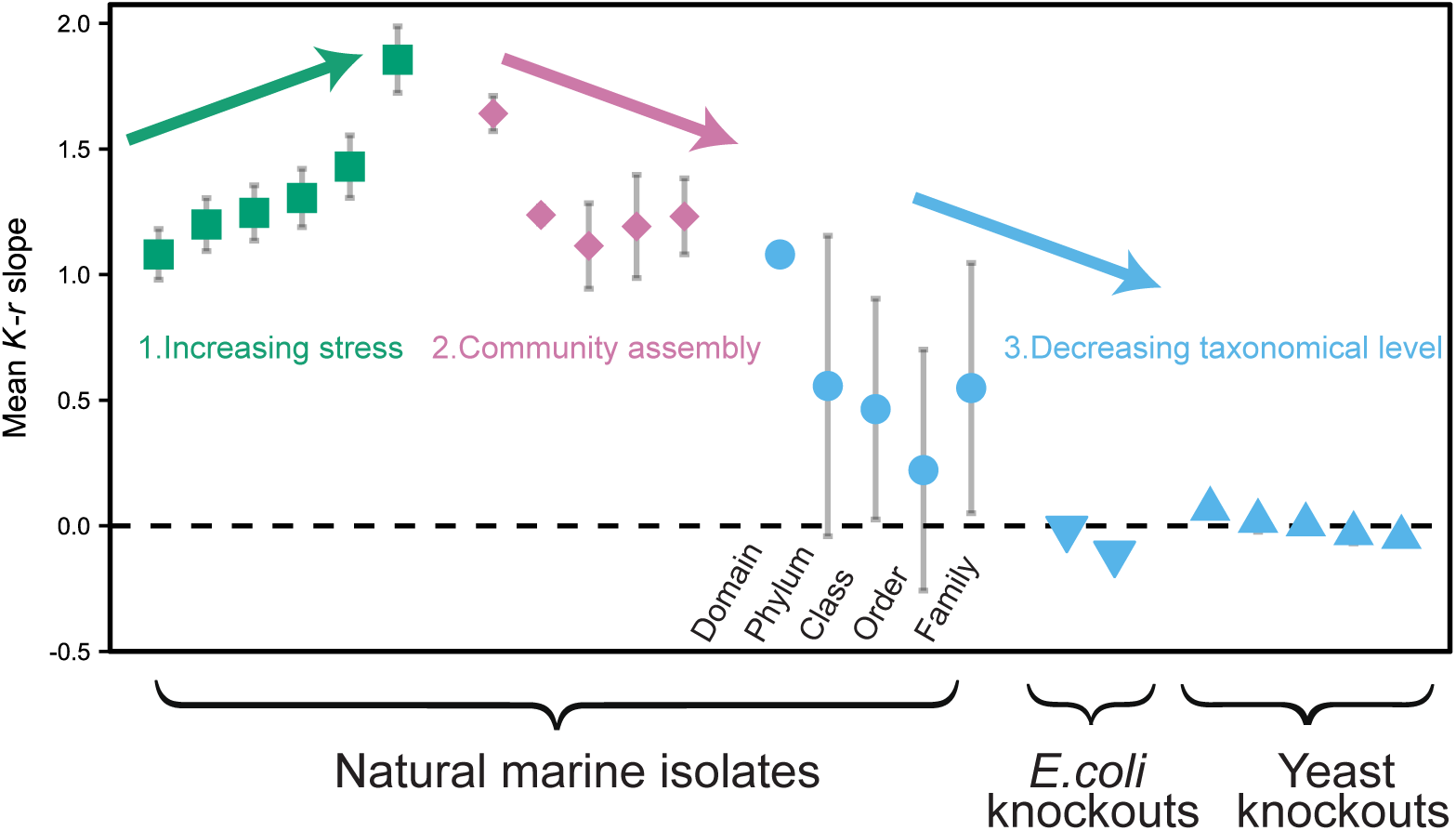
Overview of mechanisms that impact the *r*–*K* trade-up. Increasing environmental stress strengthens the positive *r*–*K* relationship (Fig. 3A), while community assembly weakens it (Fig. 4B). The trade-up also weakens at finer taxonomic resolution: both when comparing natural aquatic isolates at increasingly fine taxonomic resolution (Fig. 1F) and when analyzing single-gene knockout datasets for yeast and *E. coli* (nutrient-rich on the left, *n* = 3, 861, and nutrient-poor on the right, *n* = 3, 496). The yeast single-gene deletion dataset (*n* = 48) was analyzed under five conditions, in the following order: rich medium at elevated temperature, rich medium with added salinity stress, minimal medium, rich medium with added caffeine stress, and unstressed rich medium. Relative to unstressed rich medium, chemical stress and elevated temperature weakened the negative *r*–*K* relationship observed under standard conditions.

The distinction between within- and between-species *r*–*K* relationships has been observed in eukaryotes. In yeast, different genotypes of the same species showed trade-ups in three of nine environments and trade-offs in the other six [53], indicating that within-species *r*–*K* relationships can shift from trade-offs in high-quality environments to trade-ups under stress. This pattern is consistent with our finding that salinity stress strengthens the positive *r*-*K* relationship. Evolution experiments in *Dunaliella tertiolecta* also found that evolution under inter-specific competition increased metabolic plasticity and allowed higher *K* without reducing *r* [54]. There is also a non-significant but positive correlation between *r* and *K* in strawberry aphids [55] and a positive relationship within *Paramecium species* [56]. Although our study focuses on microbes, evidence from macro-organisms suggests that positive relationships between growth rate and carrying capacity may extend beyond microbial systems.

In the absence of an *r*–*K* trade-off, other mechanisms are necessary to explain coexistence between species. One possibility is that species are shaped by traits such as resource preference, habitat use, or temporal activity and occupy different niches [57, 11, 58]. Coexistence can arise when such niche differences outweigh the fitness differences created by a positive *r*–*K* relationship. In addition, ecological trade-offs beyond *r*–*K* may play a stabilizing role. Importantly, such trade-offs need not be present in the initial species pool, but can emerge among coexisting species through ecological filtering during community assembly. For example, models of plant competition for water predict that community assembly can select for an apparent growth rate–drought tolerance trade-off even when none exists initially [59, 60]. Thus, on top of weakening the positive *r*–*K* relationship, community assembly may simultaneously generate trade-offs along other trait dimensions, with high-*r*, high-*K* species paying costs in competitive ability, stress tolerance, or dispersal [61]. Consistent with this idea, Behringer et al. found that evolved *E. coli* under feast-and-famine cycles showed trade-offs between biofilm formation and motility, but trade-ups between growth and survival [62]. Similarly, Reding-Roman et al. found that *r* and *K* could increase together at the cost of an elongated lag phase and loss of stress-protection genes [63]. Fitness trade-offs between distinct growth phases also appear in yeast [47]. Since our findings suggest that microbial coexistence is unlikely to be explained by a simple one-dimensional *r*–*K* trade-off, a complete account of coexistence must consider trade-offs beyond *r*–*K* or other stabilizing mechanisms that allow species to coexist.

Our results suggest that *r*–*K* trade-offs are not a general feature of microbial life histories, and indeed across bacterial species growth rate and yield may often increase together. This trade-up is potentially explained by maintenance costs, but more work is required to clarify how maintenance costs vary between species and between environments. This does not mean that trade-offs are absent from microbial systems. Instead, trade-offs are likely to be found within lineages, along other trait axes, or among the subset of species that survive community assembly. Identifying which trade-offs actually structure microbial communities, and whether they are intrinsic to cell physiology or generated by community assembly itself, remains an open challenge for reconciling life-history theory with the diversity observed in nature.

## Methods

### Primary dataset: isolate collection and growth measurements

As primary dataset, we use a collection of isolates obtained from three aquatic environments [36]. In brief, isolates were collected by concentrating and plating surface water from three locations across a natural salinity gradient. The locations were designated as “Fresh-water” (the Charles River, 4 g L*^−^*^1^, 10*^◦^*C), “Estuary” (the Boston Harbor, 30 g L*^−^*^1^, 6*^◦^*C), and “Marine” (Nahant peninsula, 35 g L*^−^*^1^, 7*^◦^*C). The last group also includes isolates that were separately extracted from seaweed suspended in seawater from Nahant (called “Fucus” below). The isolates stem both from direct plating of the natural community sample (‘D0’) and plating after 2 weeks of laboratory propagation at 3 different salinities (‘D7’). Isolates were sent for 16S rRNA gene Sanger sequencing, and classified at the Genus level using the SILVA reference database (v138.1) [64].

The growth rates of these aquatic isolates were measured across 12 salinities (0, 5, 15, 20, 30, 35, 40, 45, 50, 65, 80 and 100 g L*^−^*^1^) in rich media. Nutrients were kept comparable to marine broth (MB) while the five principal sea salts (NaCl, MgCl_2_, MgSO_4_, CaCl_2_, and KCl) were varied. For each strain, independent replicate colonies were picked on at least three separate days, grown overnight, diluted 1:10000 and grown at 20 *^◦^*Cfor 48hr in a Tecan plate reader. Optical density (OD) measurements were smoothed across a 21-min window (7 data points) and the derivative of log-transformed OD was computed using the R package gcplyr, corresponding to an average growth rate over a 1-h window. The maximum growth rate at each salinity was recorded as *r*_max_. To reduce noise, the first 3 h of growth were excluded. For strains that never exceeded OD = 0.1, *r*_max_ was set to zero.

In this work, we calculated the mean growth rate (*r*) and carrying capacity (*K*) for each isolate in each condition (*n >* 3 replicates), along with their associated standard deviations. Isolates that did not grow (*r* = 0 or *K* = 0) were excluded from further analysis. We also calculated species-level averages of *r* and *K* across isolates belonging to the same species.

### *r*–*K* relationship and taxonomic variation

To examine whether the *r*–*K* relationship varies across phylogenetic scales, we first grouped isolates according to taxonomic rank (e.g., all isolates of the same family). Within each group, we estimated the slope of the *r*–*K* relationship (e.g., one slope per one family). We then averaged these slopes across groups (e.g., across all families) to obtain a representative mean slope for that taxonomic rank. This procedure was applied at the level of domain, phylum, class, order and family. Slopes were not calculated at the genus or species level because reliable estimates would require at least three isolates within each taxonomic group.

Because taxonomic groups differ widely in how many isolates they contain, we asked how sensitive the estimated *K*–*r* slope is to the number of data points used to fit it. We addressed this in two independent ways, both plotted on the same axes (slope versus number of data points). First, we plotted every individual taxonomic group as a single point, showing the slope estimated within that group against the number of isolates it contains, with error bars giving the standard error of the slope. This shows directly how estimate precision varies with group size in the real data. Second, and independently of taxonomy, we quantified the expected behaviour of the slope as a function of group size by repeated within-group subsampling. For a range of fixed group sizes *n* (*n* = 5 – 60), the isolate-level data were randomly partitioned into non-overlapping groups of size *n*, a *K*–*r* slope was fitted within each group, and the mean slope across groups was recorded, with the SEM across group slopes as its uncertainty. This partitioning was repeated 50 times per group size, each repeat contributing one point.

We next examined how the between-group *r*–*K* relationship changes with phylogeny. Using isolate-level measurements, we first calculated the mean *r* and mean *K* for each species and estimated the *K*–*r* slope across species. We then repeated this procedure at higher taxonomic levels by averaging *r* and *K* within each genus and estimating the *K*–*r* slope across genera. This approach was extended to progressively higher taxonomic ranks to evaluate how the *r*–*K* relationship varies between taxonomic groups.

To test whether the between-group slope depends on the level of aggregation itself, we repeated the analysis with taxonomy-free groups. Isolate-level data at 30 g/L salinity were randomly partitioned into non-overlapping groups of *n* isolates (*n* = 3, 6, 9, 12, 15), *r* and *K* were averaged within each group, and a single *K*–*r* slope was fitted across the group means. This was repeated for 50 independent random partitions per aggregation size, each yielding one slope estimate with its regression standard error.

### Single-carbon resource dataset

To test the generalizability of our *r*-*K* relationship and test the impact of single vs. complex carbon media, we used the microbial growth dataset from Gralka *et al.* [38]. Their complete list of strains, including taxonomy and isolation details, is provided in [38]. In brief, all experiments were conducted in liquid culture using either marine broth (MB) or a defined minimal medium with 30 mM of various carbon sources. Prior to phenotyping, marine microbes were streaked from frozen stocks and grown in rich medium for 3 days. Then, cultures were diluted 1:10 into carbon-free minimal medium for 2 h and subsequently inoculated (1:15 dilution) into minimal medium with a single carbon source. Growth was measured using optical density at 600 nm (OD_600_). Measurements were taken at least once daily (twice daily at early time points) for 17 days. Growth curves exceeding a minimum OD threshold (*>* 0.05) were fitted using a custom logistic-growth model. The fitted parameters included the maximal exponential growth rate (*r*_max_), lag time (*τ*), maximum density (*A*_1_), decay rate after peak density (*d*), and final density (*A*_2_). For each curve, three initial parameter sets were tested and the fit with the highest *R*^2^ was retained. The nonlinear fits achieved a mean *R*^2^ of 0.993 (95% CI: 0.977–0.9996). Reported *r* and *K* values correspond to the mean across all experiments in which a given strain was included.

Here, we restricted analyses to substrates with more than 50 isolates (*n* = 34) and retained only *r*, *K* estimates for isolates where the *r*–*K* relationship could be fit by linear regression with *R*^2^ *>* 0.

### Resource uptake model including maintenance cost

We developed a simple resource model that incorporates external resource uptake, intracellular resource dynamics, reproduction, and a constant metabolic maintenance cost. The model links these cellular processes to population-level parameters. This allows us to determine species-level growth rate (*r*) and carrying capacity (*K*) and explore a mechanistic explanation for the observed *r*–*K* trade-up.

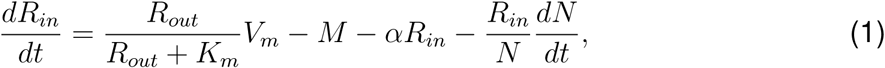

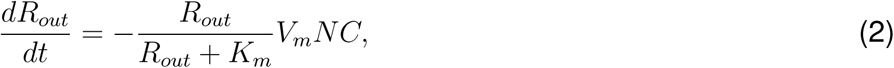

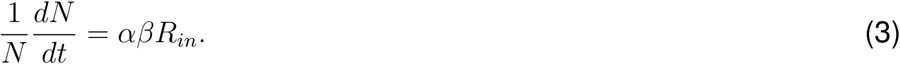

Where *R_out_*and *R_in_*describe the external and internal resource concentration respectively, and *N* the total population size. External resources are taken up by cells following a Monod-type uptake function with maximum uptake rate *V_m_* and half-saturation constant *K_m_* (i.e. the resource level at which uptake is *V_m_/*2). The uptake flux increases intracellular resources but is offset by a constant metabolic maintenance cost *M* and dilution due to cellular growth. Population growth depends on the availability of intracellular resources allocated to biomass production. Specifically, a fraction *α* of intracellular resources contributes to cell division, and the efficiency of converting intracellular resources into population growth is given by *β*. External resource depletion is scaled by a transport efficiency parameter *C*, which represents the inverse efficiency with which external resources are converted into intracellular resources (the “inverse fueling efficiency” of the cell). Together, these processes couple resource uptake, intracellular metabolism, and population growth, allowing us to investigate how a non-zero maintenance cost influences the relationship between growth rate *r* and carrying capacity *K*.

All parameters and typical simulation values are reported in Table 1. Because the model was intended to illustrate the qualitative consequences of resource uptake and maintenance rather than to represent a particular substrate or organism, resource concentrations and population sizes were expressed in arbitrary model units.

**Table 1:** Parameter values used in the resource-explicit growth model. Resource and population variables are expressed in arbitrary model units.

| Parameter | Description | Value | Units |
| --- | --- | --- | --- |
| $K_m$ | Half-saturation constant | $2.0 \times 10^9$ | resource volume<sup>-1</sup> |
| $V_m$ | Maximum uptake rate | 7 | resource (cell volume)<sup>-1</sup> h<sup>-1</sup> |
| $\alpha$ | Allocation rate of resource for division | 0.85 | h<sup>-1</sup> |
| $\beta$ | Resource to cell conversion coefficient | 0.85 | (cell volume) resource<sup>-1</sup> |
| $M$ | Maintenance cost | 0.0–4.0 | resource (cell volume)<sup>-1</sup> h<sup>-1</sup> |
| $C$ | Resource transport coefficient | 12 | cell volume population<sup>-1</sup> |
| $R_{out}$ | External resource | $2.0 \times 10^{10}$ | resource volume<sup>-1</sup> |
| $R_{in}$ | Internal resource | 0.01 | resource (cell volume)<sup>-1</sup> |
| $N$ | Population density | 10 | population volume<sup>-1</sup> |
| $t$ | Simulation time | 40 | h |

To test how maintenance costs shape the response of the *r*–*K* relationship to a change in uptake capacity, we drew a single cloud of *n* = 50 with species-specific maintenance cost *M* ∼ N (3, 0.75) and divisional resource allocation rate *α* ∼ N (0.1, 0.02). Holding *C* = 3.5, *K_m_* = 2×10^9^, *β* = 0.85 as constant. Each species was simulated twice, under high (*V_m_* = 40) and low (*V_m_* = 5) uptake rate. From each simulated growth curve we extracted *r* and *K*, mirroring how these traits are estimated from experimental data, and compared the slope of *K* ∼ *r* between conditions by linear regression. Trait-space trajectories are shown as arrows connecting each species’ initial and final (*r, K*). Within-species fractional changes *K*_final_*/K*_initial_ and *r*_final_*/r*_initial_ were plotted against *M* and compared between species with the smallest and largest 50% of *M* .

### Generalized Lotka–Volterra model and community assembly

Community dynamics were modeled using a generalized Lotka-Volterra framework with logistic growth and pairwise species interactions. The abundance *N_i_* of species *i* changes according to:

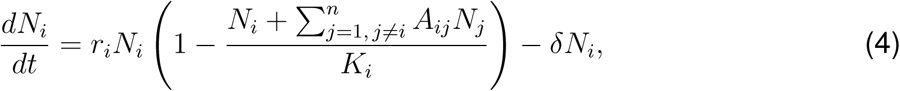

where *r_i_*is the intrinsic growth rate of species *i*, *K_i_*is its carrying capacity, and *A_ij_*represents the per-capita effect of species *j* on species *i*. The summation term captures inter-specific interactions across the community, excluding self-regulation (*A_ii_*). The parameter *δ* represents a constant mortality or dilution rate applied uniformly across species.

To simulate community assembly, we generated pools of 50 species with independently sampled intrinsic growth rates *r_i_* and carrying capacities *K_i_*. Both parameters were sampled from Gaussian distributions with mean 1.0 and standard deviation 0.2, unless otherwise stated, whereby sampled values less than or equal to 10*^−^*^6^ were rejected and resampled. Pairwise interaction coefficients were also sampled uniformly between 0 and 2*Ā*, where *Ā* denotes the mean interaction strength. Diagonal elements *A_ii_* were set to zero because intraspecific density dependence was represented separately by *K_i_*. *δ* ranges from 0 to 0.8. For each simulated community, initial abundances were sampled from a positive Gaussian distribution with mean 0.1 and standard deviation 0.02. Community dynamics were then integrated numerically using an adaptive Runge–Kutta solver over 500 generations.

At the end of each simulation, species were classified as survivors if their final abundance exceeded a threshold of 10*^−^*^4^. Species below this threshold were treated as extinct. We then compared the *r*–*K* relationship before and after community assembly. The initial relationship was calculated using all species in the starting pool, whereas the final relationship was calculated using only surviving species. For both the initial and final pools, we fit a linear regression with intercept between *r* and *K*,

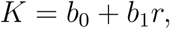

where *b*_1_ is the *K*–*r* slope.

To determine how mortality affects the emergent *r*–*K* relationship, we repeated the assembly simulations across a range of *δ* values. We generated 50 independent species pools and interaction matrices, subjected each pool to the full range of *δ* values, and then averaged the final *r*–*K* regression slopes across the 50 replicate pools for each value of *δ*. We also performed a separate sweep over mean interaction strength while fixing *δ* = 0. This control analysis tested whether increasing interaction strength alone was sufficient to change the *r*–*K* relationship.

### Community assembly data

In addition to measuring isolate growth rates, Huisman et al. also propagated natural microbial communities under laboratory conditions for 14 days [36]. The four environmental inocula (Charles labeled as Freshwater, ICA at Boston Seaport labeled as Estuary, Nahant labeled as Marine 1 and Fucus labeled as Marine 2) seeded three replicates at 3 salinities and two temperatures, yielding 18 propagated communities per inoculum (72 communities total). Cultures were transferred every 2 days at a 1:30 dilution, and the communities from day 0, 2, 6, 10, and 12 were submitted for 16S rRNA gene sequencing. Amplicon sequence variants (ASVs) were identified using DADA2 [65] and assigned species identity using the SILVA database (v138.1)[64]. To correct relative abundances for variation in 16S rRNA gene copy number, copy numbers were obtained from rrnDB and assigned to each ASV based on the lowest taxonomic level available. In addition, ASVs were matched to isolate 16S rRNA gene sequences using BLAST, requiring *>* 97% sequence identity [66].

To quantify the *r*–*K* relationship, each ASV was matched to its closest isolate, and the corresponding estimates of *r* and *K* were used in subsequent species-level analyses. Because low-abundance ASVs are less reliably quantified and are represented by relatively few isolates, we restricted the analysis to ASVs with a relative abundance greater than 0.1% in each community at each time point. Sensitivity analyses using more relaxed (*>* 0.05%) and more strict (*>* 0.15%) thresholds yielded qualitatively similar results (Extended Data Fig. 7).

### Experimental *r*–*K* relationship in the context of controlled gene knockouts

To test the *r*–*K* relationship within strains, we analyzed published single-gene knockout datasets from *Escherichia coli* [50] and *Saccharomyces cerevisiae* [52]. The *E. coli* dataset measured growth curves for single-gene knockout strains from the Keio collection in rich (LB) and minimal medium (M63) liquid culture. We used the growth rate *r* and carrying capacity *K* values reported in the original study, but filtered to retain only data with the highest validation indicator for both *r* and *K*, corresponding to measurements without evidence of computational bias in *r*, biological bias in *K*, or replicate outliers. We then averaged *r* and *K* across biological replicates (*n* = 3) for each genotype-by-medium combination. Cases with *r* = 0 or *K* = 0 were removed. We examined the *r*–*K* relationship by fitting ordinary least-squares linear regressions of carrying capacity against growth rate, *K* ∼ *r*, separately for each medium and gene knockout class. Following the convention of [50], we excluded genes with partial or unknown functions and grouped predicted functional categories with their corresponding annotated categories.

For the yeast analysis, we used the genetic perturbation dataset from [52]. In this study, the authors carried out 96 crosses of haploid yeast strains, each carrying a different single-gene deletion. We refer to these as deletion A and deletion B within each cross. Each cross produced four haploid daughter genotypes: a strain with no deletion (‘wild-type’), one strain each carrying either deletion A or B only (‘single-deletion’), and a strain carrying both deletion A and B (‘double-deletion’). This produced 384 strains in total. The 96 wild-type yeast strains, 192 single-deletion strains, and 96 double-deletion strains were each grown in monoculture in rich medium at 30*^◦^*C, with yield and maximum intrinsic growth rate measured for each strain. In addition, Jakubowska et al. cultured a subset of 48 single-deletion strains in five environments: rich nutrient medium (YPD broth), poor nutrient medium (SD defined medium), rich medium supplemented with caffeine, rich medium supplemented with NaCl, and rich medium at elevated temperature. All environments were assayed at 30°C, except for the elevated-temperature treatment, which was assayed at 37°C. Each genotype-by-environment combination had two biological replicates. We used dry mass per glucose consumed as an approximation of carrying capacity (since it reflects biomass yield). We removed entries with missing values and fit a linear regression between maximum growth rate and dry mass per glucose consumed. We used this to test whether genetic perturbations in yeast produced a positive, negative, or non-significant *r*–*K* relationship.

## Data availability

Upon publication, all data and code required to reproduce the analyses and figures will be made publicly available in a permanent repository.

## Acknowledgements

The authors thank the Gore lab for helpful discussions about this work. In addition, we thank Benjamin Gore and Martina Dal Bello. JSH was supported by Human Frontier Science Program (HFSP) Postdoctoral Fellowship LT0045/2023-L (DOI: 10.52044/HFSP.LT00452023-L. pc.gr.171493). ZX acknowledges partial support from the Thomas Frank Fellowship from MIT Physics Department.

## Author contributions

Z.X., J.S.H. and J.G. conceptualized the study. Z.X. and J.S.H. carried out formal analysis, investigation, and wrote the original draft. Z.X., J.S.H. and J.G. jointly contributed to subsequent writing, review and editing. Supervision was carried out by J.S.H. and J.G.

## S1 Extended Data Figures

**Extended Data Fig. 1.**
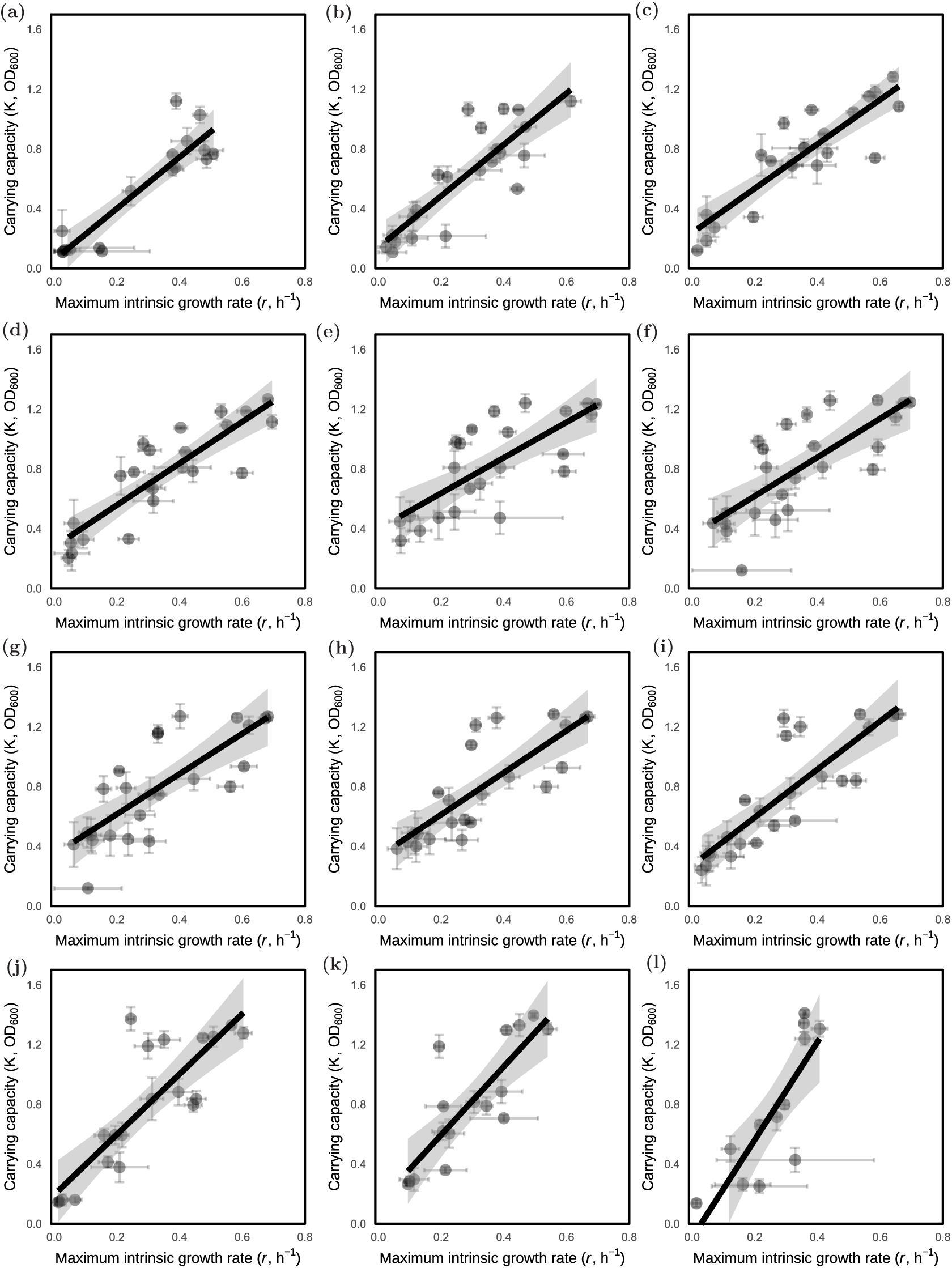
The *r*–*K* relationship remains positive across salinities. **(a–l)** Relationships between maximum intrinsic growth rate, *r*, and carrying capacity, *K*, across species grown at salinities of 0, 5, 15, 20, 30, 35, 40, 45, 50, 65, 80, and 100 g L*^−^*^1^ (left to right, top to bottom). Each point represents the mean of a species, with error bars indicating variation across isolates of the same species (SEM). Lines show linear regressions calculated at each salinity, and gray shading indicates the 95% confidence intervals. Isolates that did not grow at a particular salinity (*r* = 0, *K* = 0) were excluded after calculating species-level values by averaging, and before subsequent regression analyses.

**Extended Data Fig. 2.**
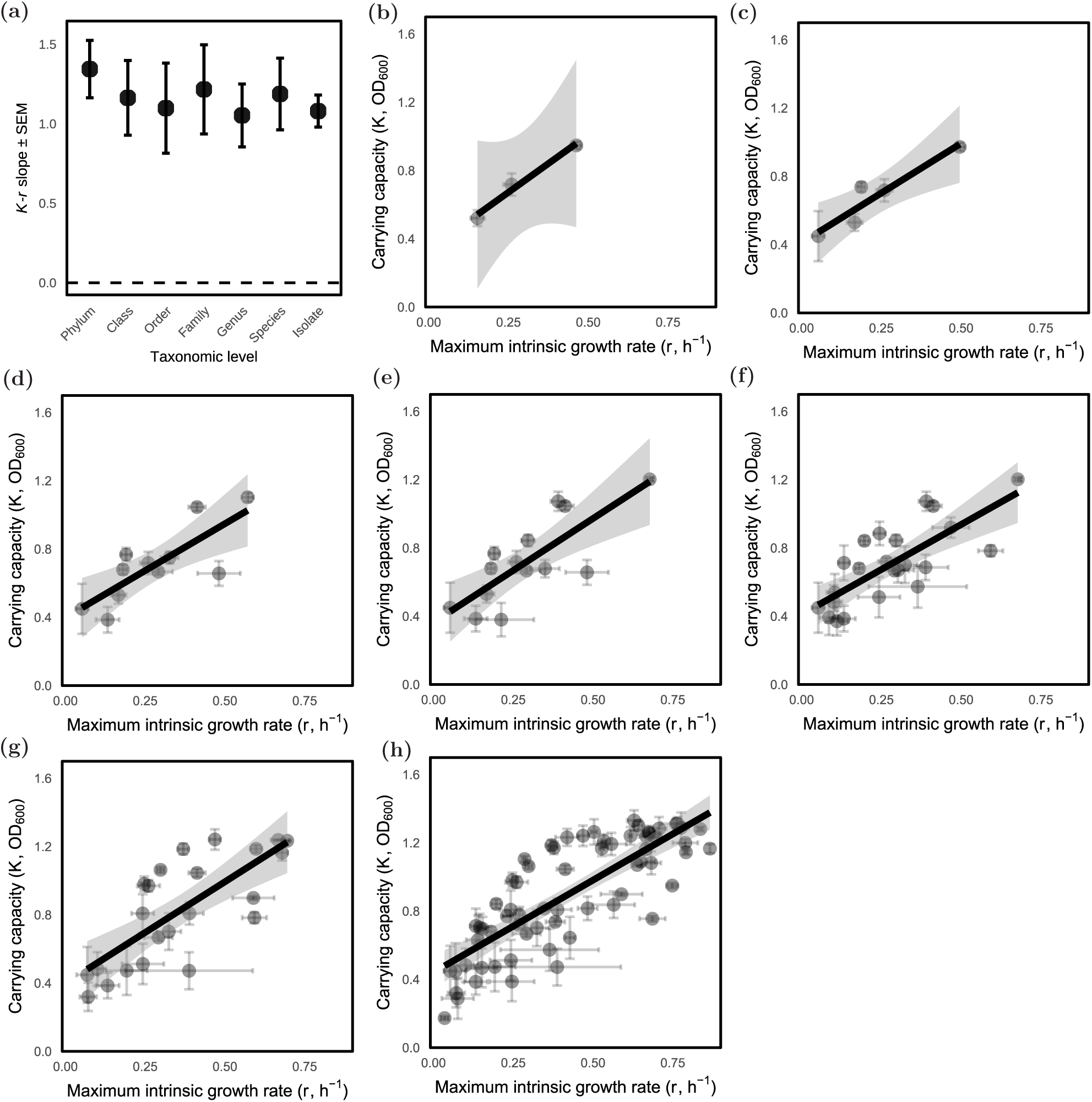
The *r*–*K* trade-up remains consistently positive after aggregating isolates at different phylogenetic levels. **(a)** Regression slopes between maximum intrinsic growth rate *r* and carrying capacity *K*, after averaging at different taxonomic levels. Points show the mean slope at each aggregation level, and error bars indicate the standard error of the slope. The numbers of points included in the regression are *n* = 3, 5, 11, 14, 23, 23, 66 when averaged at the phyla, class, order, family, genus, species, or isolate level respectively. The dashed horizontal line marks a slope of zero. **(b–h)** Linear regressions after aggregating data at the phylum (b; slope = 1.35, SE = 0.18, *R*^2^ = 0.98, *n* = 3), class (c; slope = 1.16, SE = 0.24, *R*^2^ = 0.89, *n* = 5), order (d; slope = 1.10, SE = 0.28, *R*^2^ = 0.63, *n* = 11), family (e; slope = 1.22, SE = 0.28, *R*^2^ = 0.61, *n* = 14), genus (f; slope = 1.05, SE = 0.20, *R*^2^ = 0.58, *n* = 23), species (g; slope = 1.19, SE = 0.23, *R*^2^ = 0.57, *n* = 23) and isolate(h; slope = 1.08, SE = 0.10, *R*^2^ = 0.64, *n* = 66) levels. Each point represents the mean *r* and *K* for one phylogenetic group, with error bars showing variation (SEM) within that group. Black lines show linear regressions, and gray shading indicates the 95% confidence interval.

**Extended Data Fig. 3.**
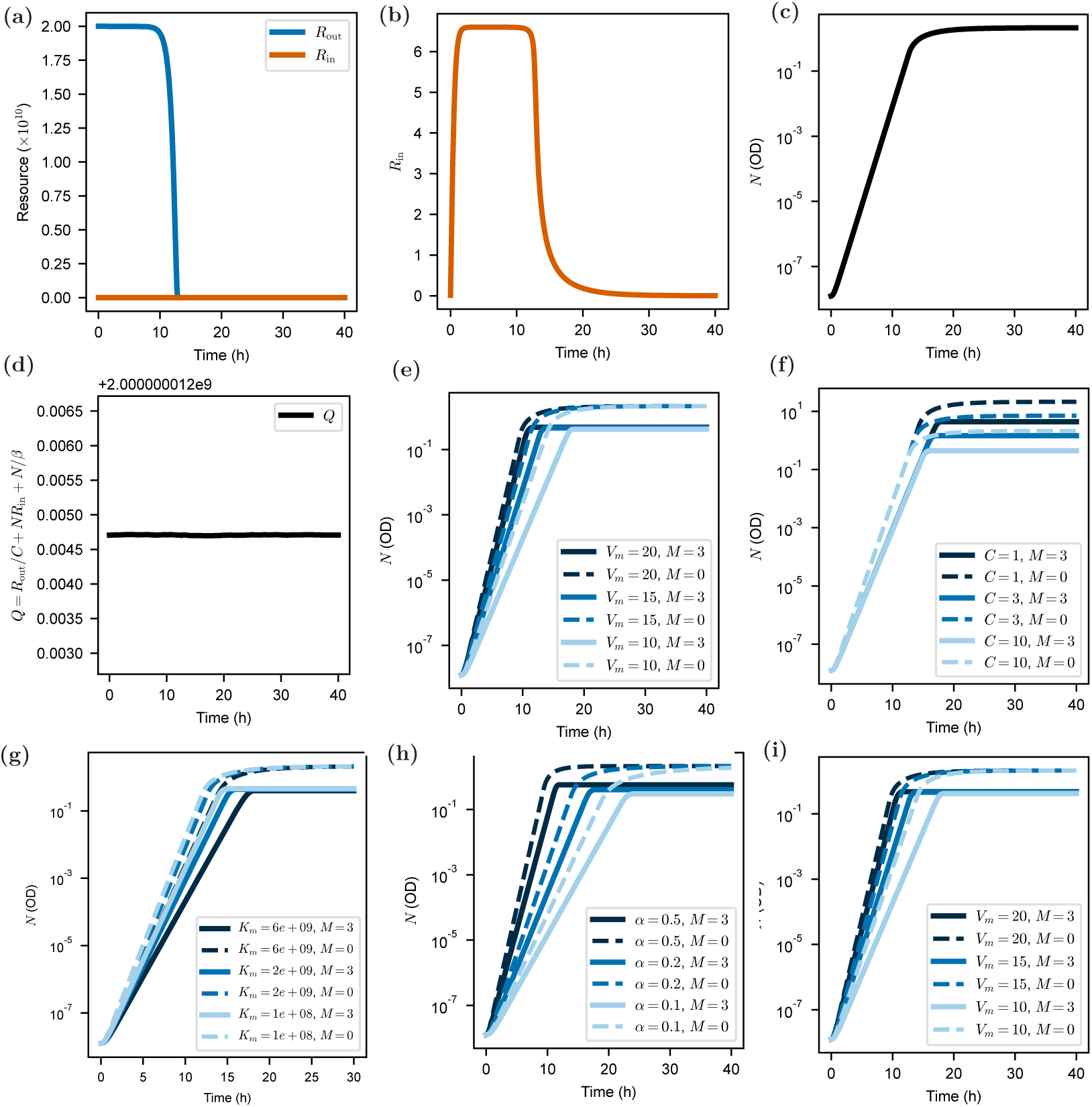
Resource-explicit growth model links physiological parameters to growth rate and carrying capacity. **(a–d)** Representative model dynamics in the absence of maintenance cost (*M* = 0). **(a)** External resource *R_out_*(blue) is consumed over time, while intracellular resource *R_in_* (orange) remains small relative to the external resource pool. **(b)** *R_in_* rapidly equilibrates before declining as external resources are depleted. **(c)** The model reproduces logistic growth of the population *N* . **(d)** Total resource and biomass are conserved, as shown by the approximately constant quantity *Q* = *R*_out_*/C* + *NR*_in_ + *N/β*. **(e–i)** Effects of individual model parameters on population growth. Solid lines represent growth with maintenance cost (*M* ≠ 0), whereas dashed lines represent growth without maintenance cost (*M* = 0). **(e)** In the absence of maintenance cost (*M* = 0), increasing the maximum uptake rate *V_m_* increases *r* without changing *K*. With maintenance cost, increasing *V_m_* increases both *r* and *K*. **(f)** Increasing the cellular “inverse fueling efficiency” *C*, which controls the conversion of external into intracellular resources, decreases *K* without changing *r*, both with and without maintenance cost. **(g)** In the absence of maintenance cost, decreasing the half-saturation constant *K_m_* increases *r* without changing *K*. With maintenance cost, decreasing *K_m_* increases both *r* and *K*, with a stronger effect on *r*. **(h)** In the absence of maintenance cost, increasing *α*, the fraction of intracellular resources allocated to cellular growth, increases *r* without changing *K*. With maintenance cost, increasing *α* increases both *r* and *K*. **(i)** Increasing *β*, the conversion efficiency of intracellular resources into population growth, increases both *r* and *K*, with or without maintenance cost.

**Extended Data Fig. 4.**
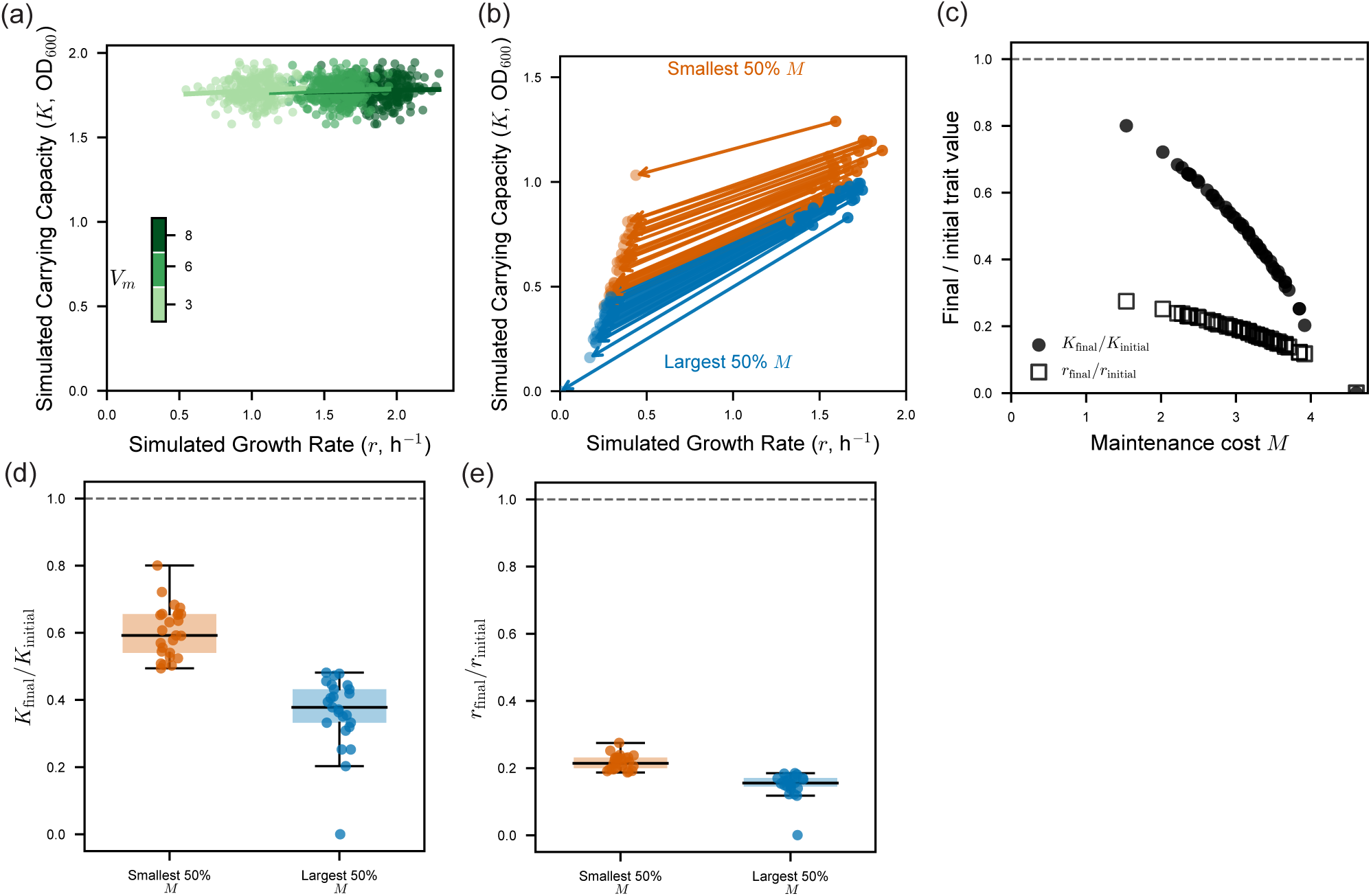
The resource model predicts that species with higher maintenance costs experience greater reductions in *K* than in *r* as *V_m_* decreases. **(a)** In the absence of maintenance costs, reducing the maximum uptake rate, *V_m_*, changes the distribution of *r* and *K* but not the slope of their relationship. **(b)** Trajectories in *r*–*K* trait space following a reduction in *V_m_*in the presence of maintenance costs. Each arrow connects the initial and final trait values. Blue trajectories represent species with larger maintenance costs *M* whereas orange trajectories represent species with smaller values. **(c)** Relative changes in *K* and *r* as a function of maintenance cost and changing *V_m_*. As *M* increases, reduced *V_m_*has a stronger effect on *K*_final_*/K*_initial_ than on *r*_final_*/r*_initial_. **(d)** Species with larger maintenance costs undergo greater fractional reductions in *K* than species with smaller maintenance costs. **(e)** In contrast, the fractional reduction in growth rate is similar between the larger- and smaller-*M* groups.

**Extended Data Fig. 5.**
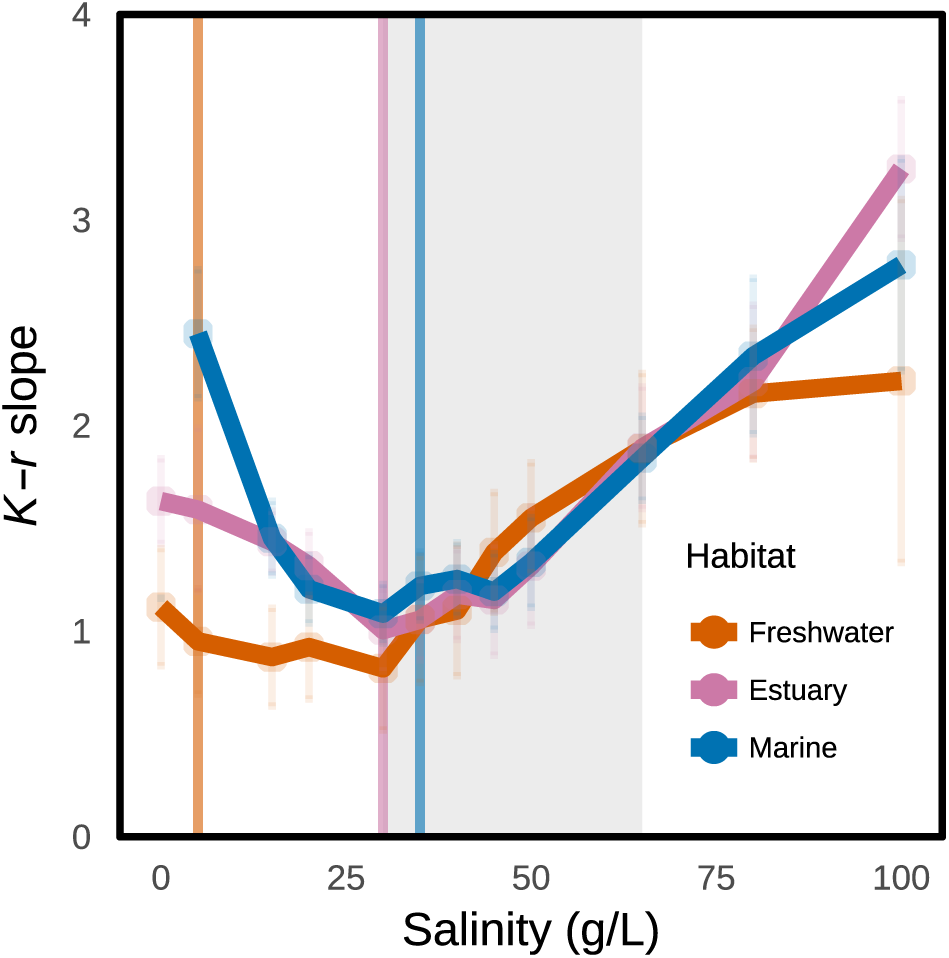
Salinity stress strengthens the positive *r*–*K* relationship away from habitat-specific optima. The slope of the relationship between maximum intrinsic growth rate *r* and carrying capacity *K* (points ± standard error) calculated at each salinity for species from each habitat. For each habitat, the slope reaches its lowest value near the apparent optimum salinity, producing a dip in the *r*–*K* slope. Away from this optimum, the slope increases. The gray shaded region marks the salinity range used in the Figure 3.

**Extended Data Fig. 6.**
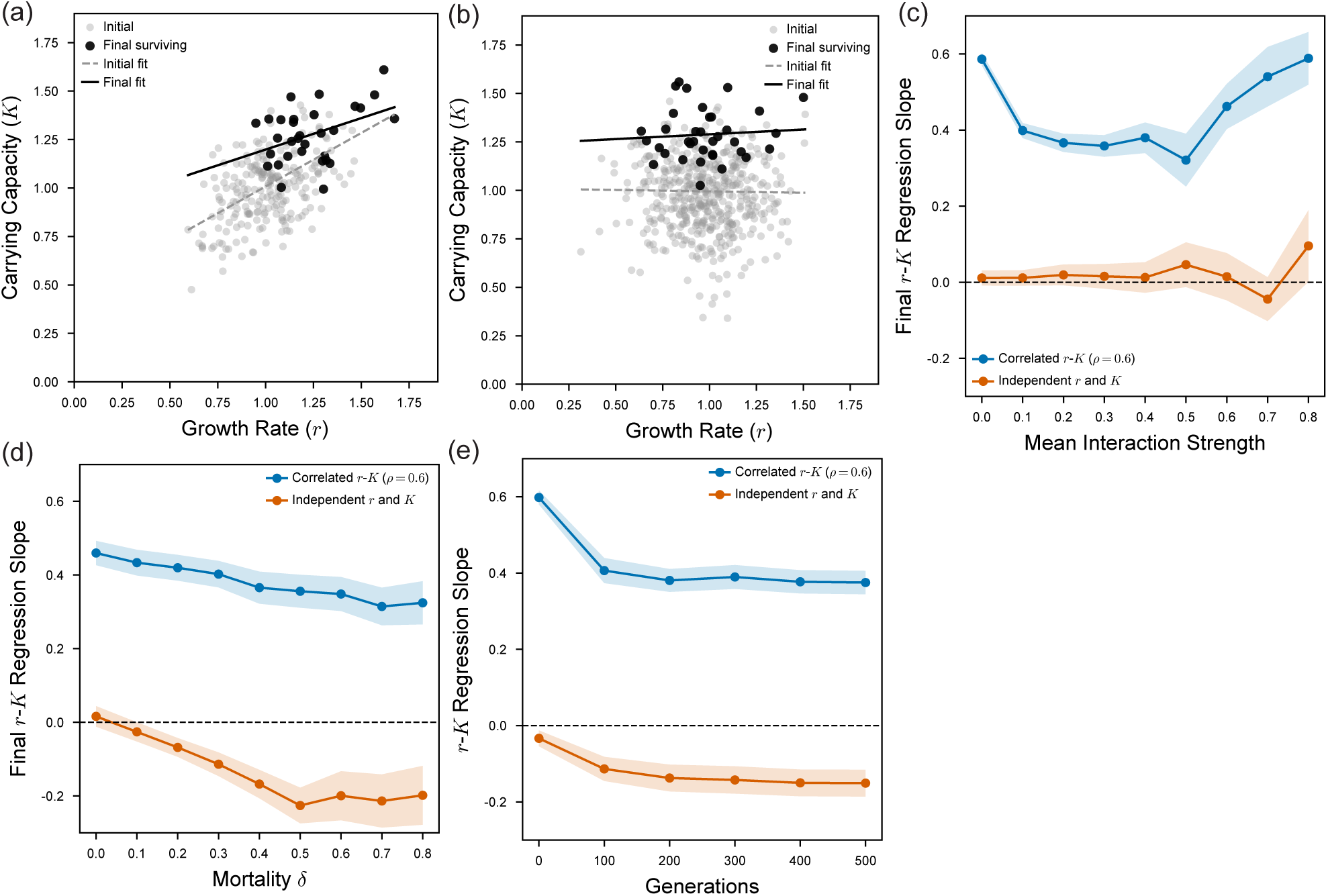
A generalized Lotka–Volterra model predicts that community assembly weakens positive *r*–*K* relationships under mortality. **(a)** Example of community assembly from an initial species pool in which *r* and *K* were jointly sampled from a correlated Gaussian distribution with correlation *ρ_rK_* = 0.6. Gray points represent the initial species pool, and black points represent species that survive after community assembly. The dashed gray line shows the initial linear regression, and the solid black line shows the regression among surviving species. **(b)** Example of community assembly from an initial species pool in which *r* and *K* were jointly sampled from an uncorrelated Gaussian distribution. **(c)** Effect of mean interaction strength on the final *K*–*r* regression slope among surviving species for initially correlated and uncorrelated *r* and *K*. Increasing mean interaction strength does not systematically affect the final slope, which remains close to zero across interaction strengths. **(d)** Effect of mortality, *δ*, on the final *K*–*r* regression slope among surviving species for initially correlated and uncorrelated *r* and *K*. Increasing mortality progressively reduces the final slope. **(e)** Change in the *r*–*K* regression slope during community assembly for initially correlated and uncorrelated *r* and *K*. Starting from an initially positive relationship (slope = 0.6), the slope decreases over successive generations under mortality (*δ* = 0.3) but remains positive.

**Extended Data Fig. 7.**
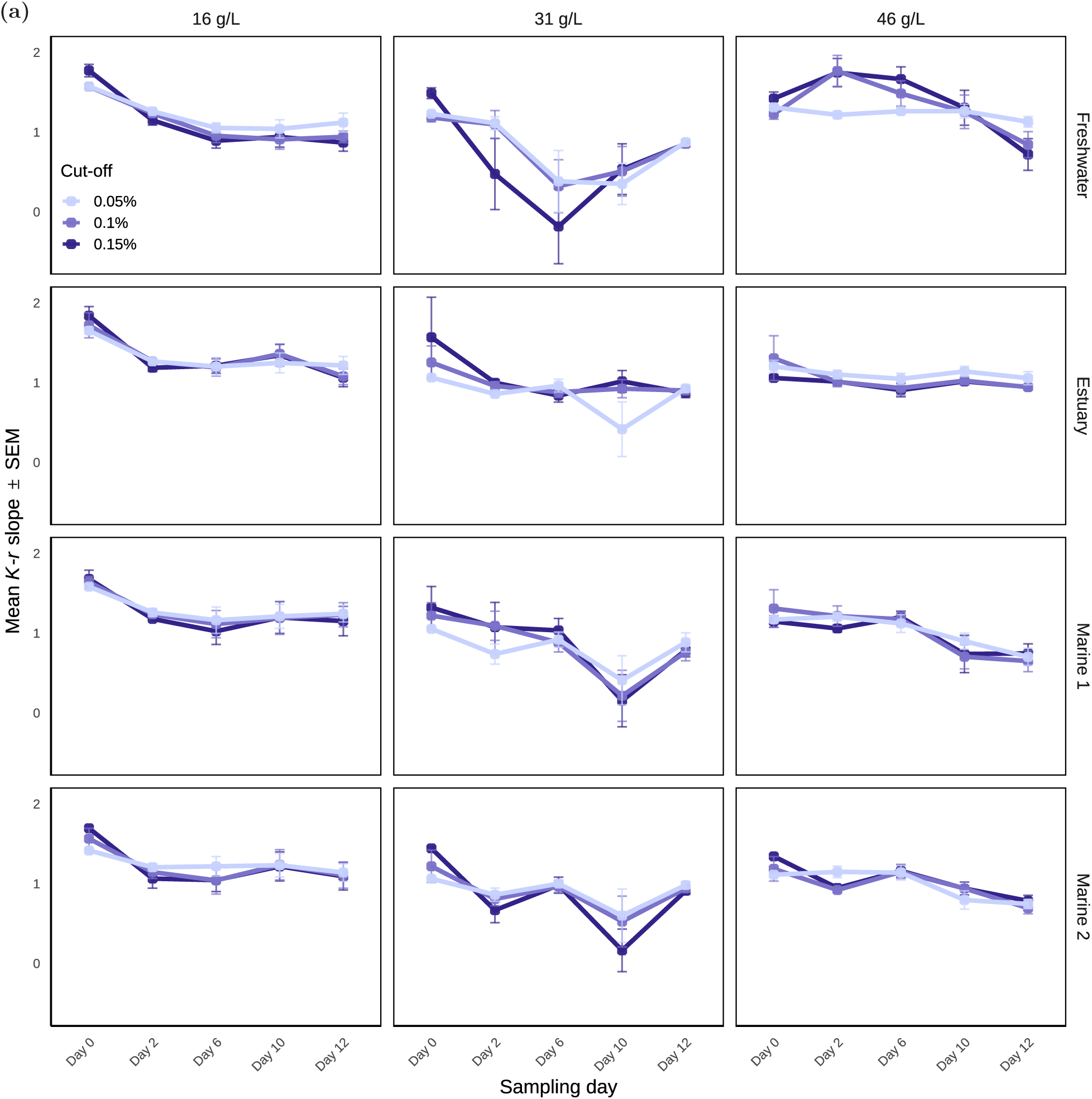
Experimental community assembly weakens the positive *r*–*K* relationship over time. Mean *K*–*r* regression slope ± SEM across sampling days during community assembly. Panels show communities from different inoculum habitats and salinity treatments. Colors indicate the relative-abundance thresholds used to determine which taxa were included in the analysis: 0.05%, 0.1%, and 0.15%. Across habitats, salinities, and relative-abundance thresholds, the *K*–*r* slope generally decreases from Day 0 to later time points.

**Extended Data Fig. 8.**
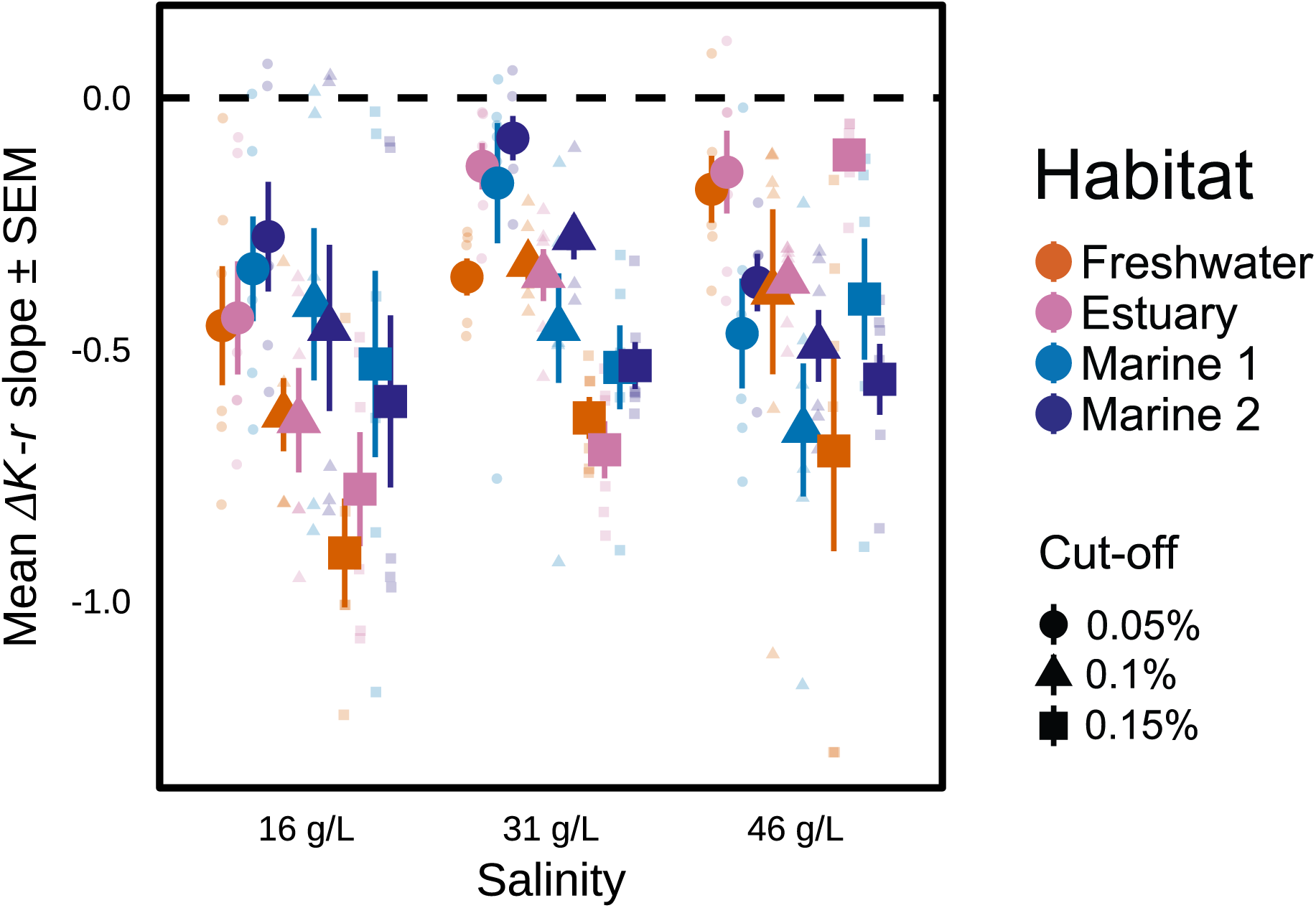
The *r*–*K* relationship weakens between the beginning and end of community assembly. Change in the *K*–*r* regression slope between the initial and final community assembly time points, calculated as Δslope = slope ∗ final − slope ∗ initial. Points show the mean Δslope± SEM for each inoculum source (colors), salinity treatment, and relative-abundance threshold (shapes: circles, 0.05%; triangles, 0.1%; and squares, 0.15%). Small, high transparency points show slope changes for individual communities. Negative values indicate that the *K*–*r* slope decreased during community assembly.

**Extended Data Fig. 9.**
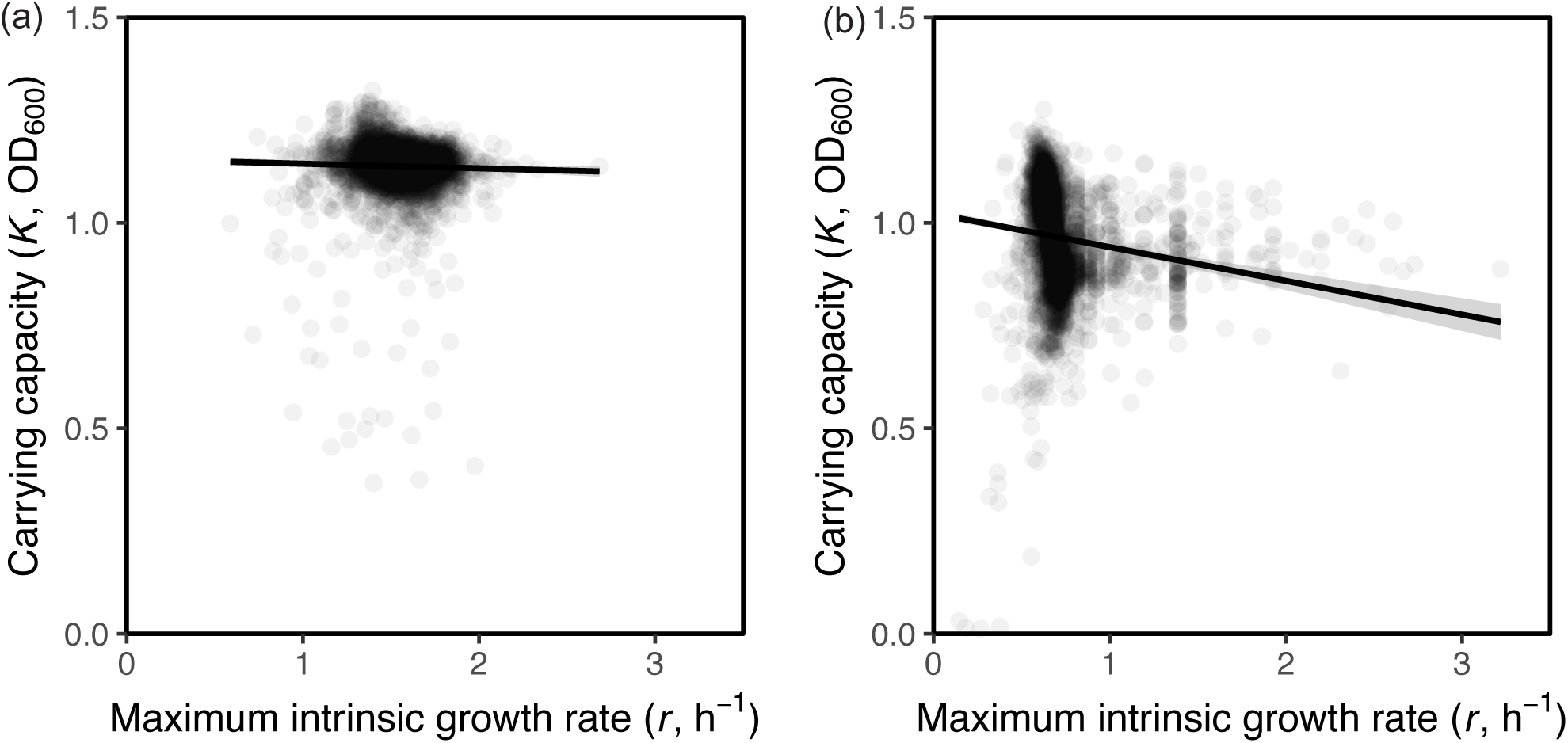
*E. coli* knockouts show no strong *r*–*K* trade-off. **(a)** Rich nutrient medium: *K*–*r* slope for *r >* 0, *K >* 0.1 is -0.01 ±0.01 (slope ± SE, *n* = 3, 861) **(b)** Low nutrient medium: *K*–*r* slope for *r >* 0, *K >* 0.1 is -0.11 ±0.01 (slope ± SE, *n* = 3, 496).

**Extended Data Fig. 10.**
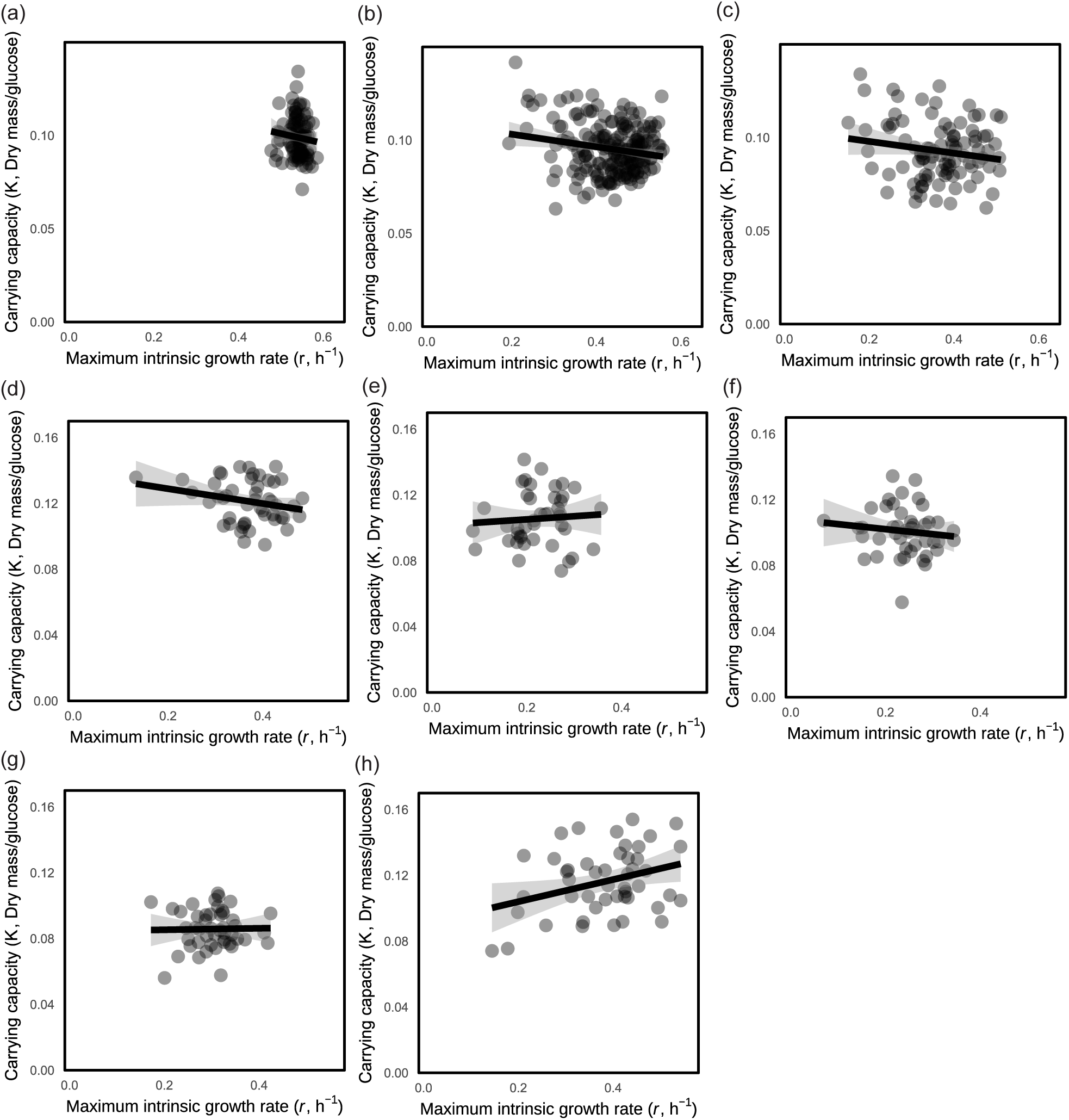
Yeast genetic perturbations show weak *r*–*K* trade-offs or no relationship. **(a)** Wild-type yeast in rich media (*n* = 96). **(b)** Single-mutant yeast in rich media (*n* = 192). **(c)** Double-mutant yeast in rich media (*n* = 96). **(d)** The subset of single-mutant yeast in rich media (*n* = 48). **(e)** The subset of single-mutant yeast in rich media supplemented with NaCl (*n* = 48). **(f)** The subset of single-mutant yeast in rich media supplemented with caffeine (*n* = 48). **(g)** The subset of single-mutant yeast in minimal media (*n* = 48). **(h)** The subset of single-mutant yeast in rich media at elevated culturing temperature (*n* = 48).

## Supplementary Figures

**Supplementary Fig. 1.**
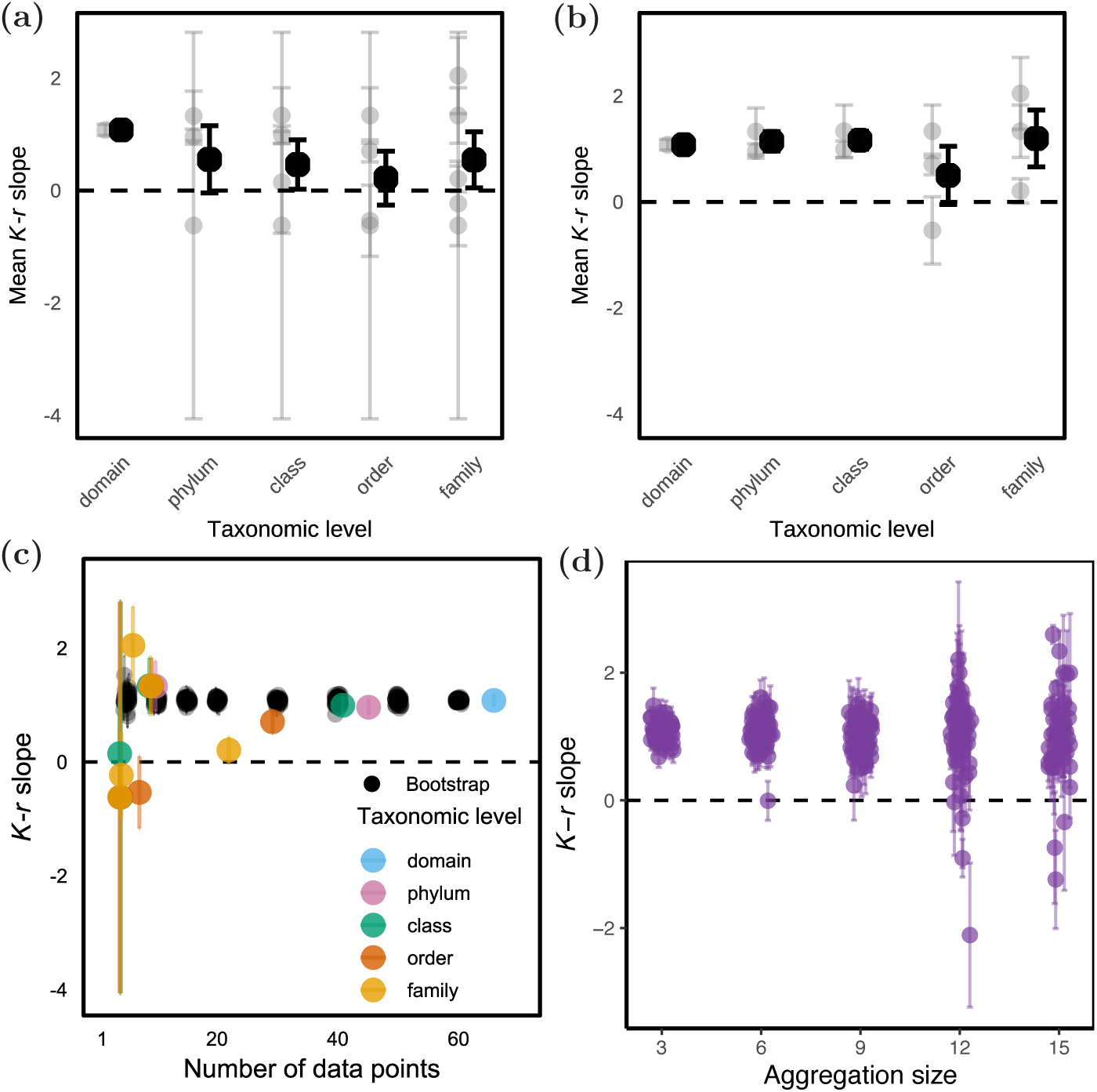
Sample size does not explain variation in the *r*–*K* relationship across phylogenetic groups. **(a)** Mean within-group *K*–*r* slope across taxonomic levels (phyla, classes, orders, or families) with *n* ≥ 3 isolates required to calculate each within-group slope. Grey points indicate *K*–*r* slopes within each individual taxonomic-group and SE estimates (*n* = 1 for domain, *n* = 3 for phylum, *n* = 4 for class, *n* = 5 for order, and *n* = 7 for family). Black points show the mean slope ± SEM at each taxonomic level. The dashed line indicates a slope of zero. **(b)** Similar to (a) but with *n* ≥ 5 isolates required to calculate each within-group slope. The number of grey points is *n* = 1 for domain, *n* = 2 for phylum, *n* = 2 for class, *n* = 3 for order, and *n* = 3 for family. **(c)** Relationship between the number of data points used to estimate each *r*–*K* slope and the resulting slope across taxonomic levels. Each colored point represents one taxonomic group, colored by taxonomic level, with error bars indicating the standard error of the slope. Black points show results from repeated within-group subsampling: for each group size, isolates were randomly partitioned into non-overlapping groups, an *r*–*K* slope was calculated within each group, and the mean slope across groups was recorded. This procedure was repeated 50 times for each group size, with each black point representing one repeat. Error bars indicate the SEM across slopes from the groups within that repeat. The dashed horizontal line marks a slope of zero. **(c)** Repeated subsampling analysis of the effect of aggregation size on the *r*–*K* slope. For each aggregation size, *n*, isolates were randomly partitioned into non-overlapping groups of *n* isolates, and *r* and *K* were averaged within each group. A single *r*–*K* slope was then fitted across the group means. This procedure was repeated 50 times for each value of *n*.

**Supplementary Fig. 2.**
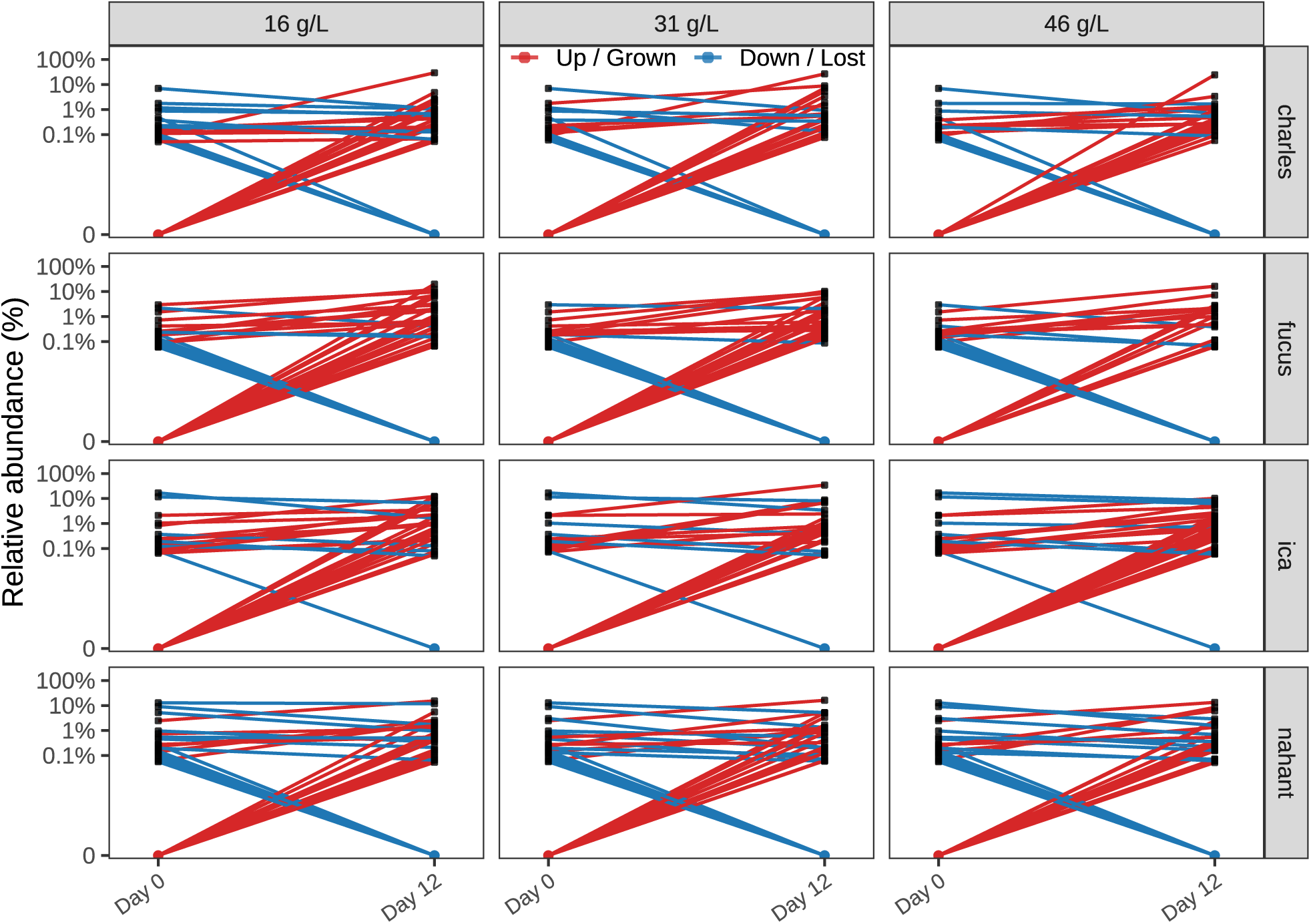
Isolate-level abundance shifts during community assembly. Each panel shows the relative abundance of individual isolates between the initial community composition at Day 0 and the assembled community at Day 12, faceted by inoculum habitat source and salinity treatment condition. Lines connect the same isolates across timepoints. Red lines indicate taxa that increased in relative abundance, whereas blue lines indicate taxa that decreased in relative abundance or were lost by Day 12. Points plotted at the bottom of the log-scaled axis represent taxa below the detection threshold: red lines originating from this point indicate isolates that were near-zero or undetected at Day 0 but increased by Day 12, while blue lines ending near-0 indicate taxa that declined to near-zero or were lost by Day 12. This visualization highlights widespread compositional turnover during community assembly, including both changes in abundance among persisting isolates and the gain or loss of detectable isolates across habitats and salinity treatments.

**Supplementary Fig. 3.**
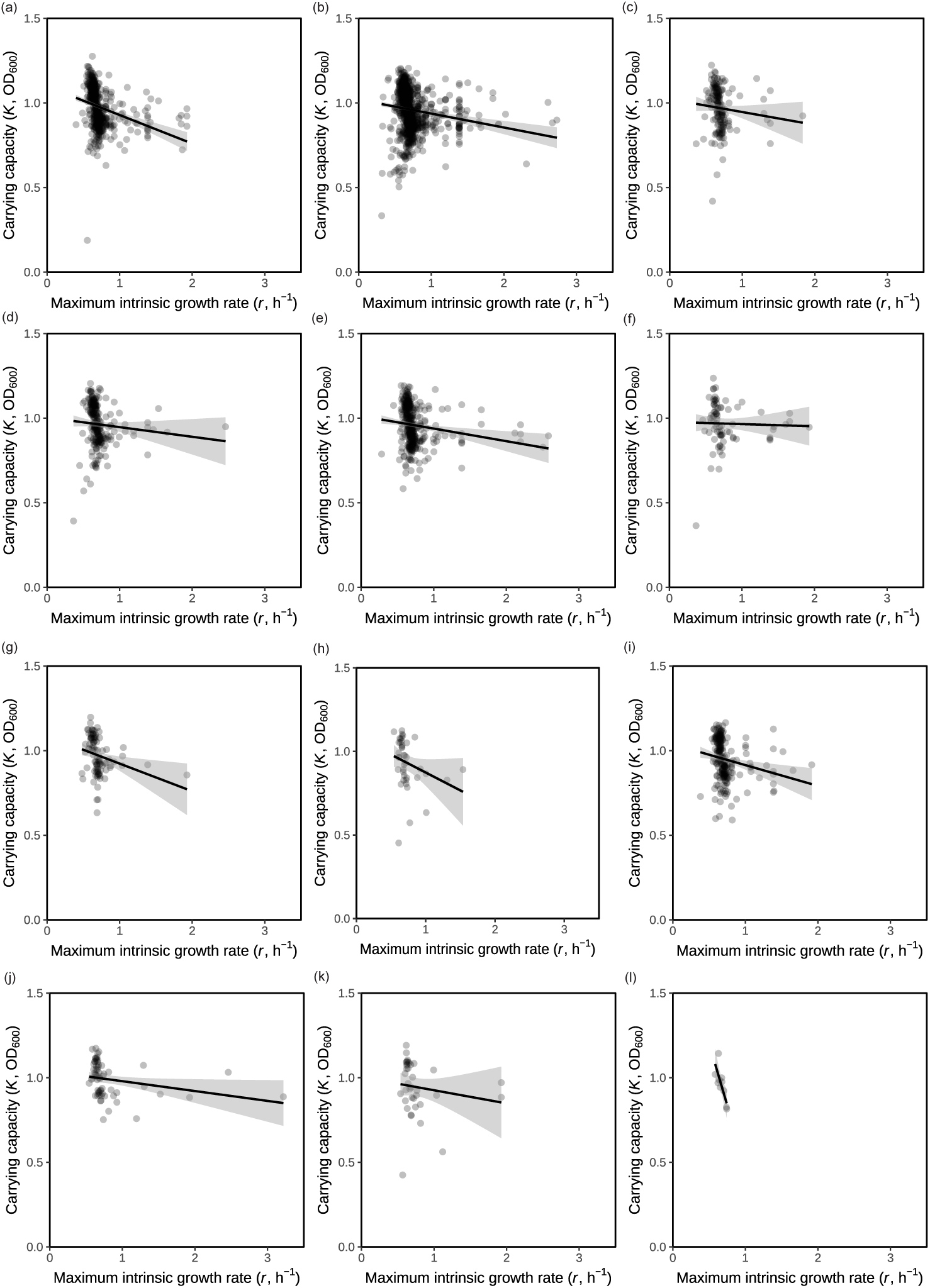
*E. coli* knockouts in minimal medium grouped by gene category display weak *r*-*K* trade-offs. Panels are ordered from left to right and then from top to bottom according to the following gene-categories: transporter (a), enzyme (b), factor (c), membrane protein(d), regulator (e), structural components (f), carrier (g), lipoprotein (h), phage/insertion sequences (IS) in common with ancestral strain (i), pseudogenes in common with ancestral strain (j), cell process (k), and leader peptide (l), respectively.

**Supplementary Fig. 4.**
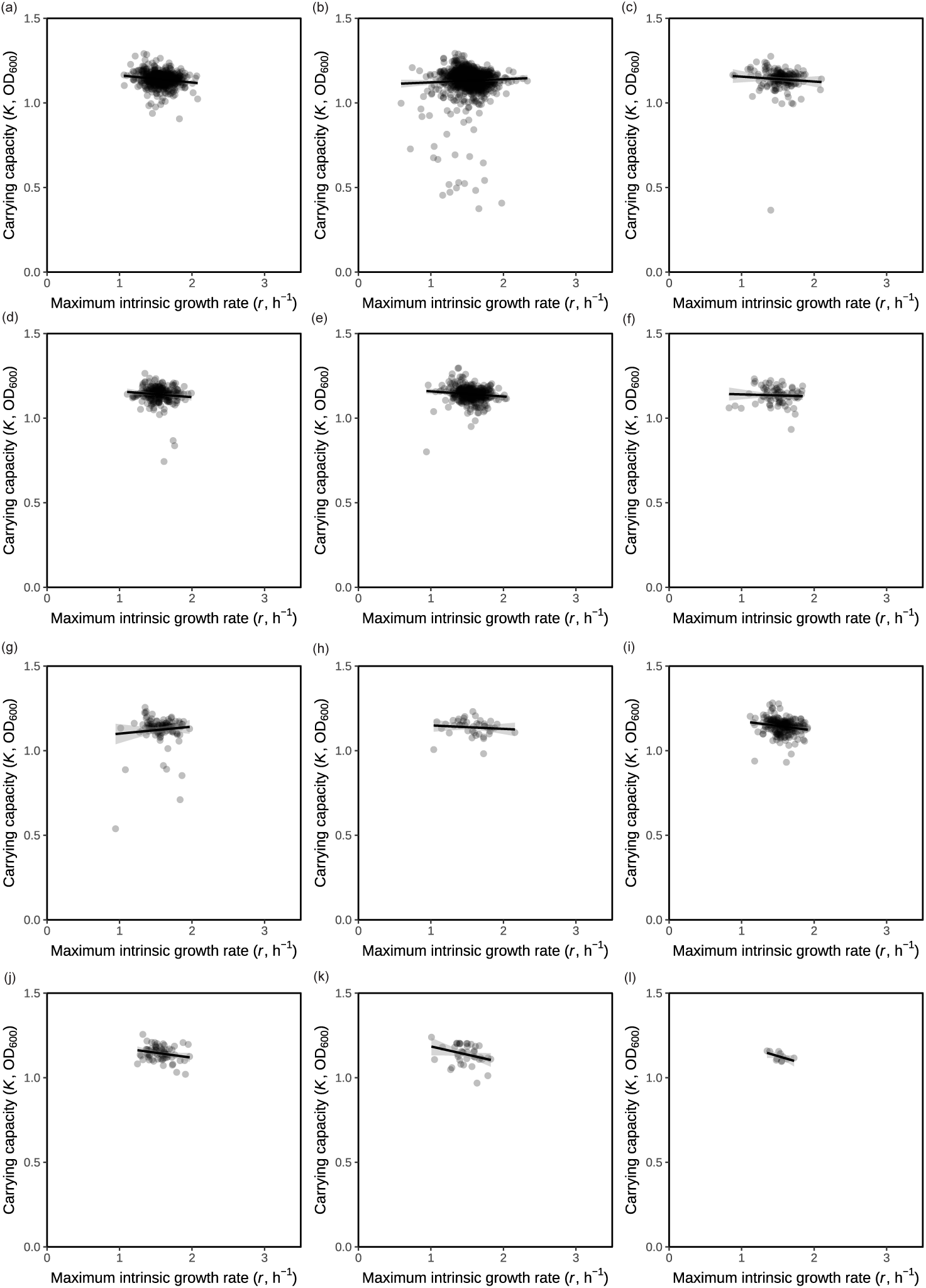
*E. coli* knockouts in rich medium grouped by gene category display weak or no *r*-*K* trade-offs. Panels are ordered from left to right and then from top to bottom according to the following gene-categories: transporter (a), enzyme (b), factor (c), membrane protein(d), regulator (e), structural components (f), carrier (g), lipoprotein (h), phage/insertion sequences (IS) in common with ancestral strain (i), pseudogenes in common with ancestral strain (j), cell process (k), and leader peptide (l), respectively.

## References

[1] Jamie M Kneitel and Jonathan M Chase. Trade-offs in community ecology: linking spatial scales and species coexistence. Ecology letters, 7(1):69–80, 2004.

[2] Eric R Pianka. On r-and k-selection. The american naturalist, 104(940):592–597, 1970.

[3] Alita R Burmeister and Paul E Turner. Trading-off and trading-up in the world of bacteria–phage evolution. Current Biology, 30(19):R1120–R1124, 2020.

[4] Thomas Alerstam, Anders Hedenström, and Susanne Åkesson. Long-distance migration: evolution and determinants. Oikos, 103(2):247–260, 2003.

[5] Rahul Bale, Max Hao, Amneet Pal Singh Bhalla, and Neelesh A Patankar. Energy efficiency and allometry of movement of swimming and flying animals. Proceedings of the National Academy of Sciences, 111(21):7517–7521, 2014.

[6] David Lack. The significance of clutch-size. Ibis, 89(2):302–352, 1947.

[7] Juan Bueno and Angel Lopez-Urrutia. The offspring-development-time/offspring-number trade-off. The American Naturalist, 179(6):E196–E203, 2012.

[8] Ivana Gudelj, Robert E Beardmore, SS Arkin, and R Craig MacLean. Constraints on microbial metabolism drive evolutionary diversification in homogeneous environments. Journal of evolutionary biology, 20(5):1882–1889, 2007.

[9] David Reznick, Michael J Bryant, and Farrah Bashey. r-and k-selection revisited: the role of population regulation in life-history evolution. Ecology, 83(6):1509–1520, 2002.

[10] Peter Chesson. Mechanisms of maintenance of species diversity. Annual review of Ecology and Systematics, 31(1):343–366, 2000.

[11] David Tilman. Resource competition and community structure. Number 17. Princeton university press, 1982.

[12] Madhav Gadgil and Otto T Solbrig. The concept of r-and k-selection: evidence from wild flowers and some theoretical considerations. The American Naturalist, 106(947):14–31, 1972.

[13] John H Andrews and Robin F Harris. r-and k-selection and microbial ecology. In Advances in microbial ecology, pages 99–147. Springer, 1986.

[14] David A Lipson, Russell K Monson, Steven K Schmidt, and Michael N Weintraub. The trade-off between growth rate and yield in microbial communities and the consequences for under-snow soil respiration in a high elevation coniferous forest. Biogeochemistry, 95(1):23–35, 2009.

[15] Peter Koefoed Bjørnsen. Bacterioplankton growth yield in continuous seawater cultures. Marine Ecology Progress Series, 30:191–196, 1986.

[16] Klaus Kristiansen, Helle Nielsen, Bo Riemann, and Jed A Fuhrman. Growth efficiencies of freshwater bacterioplankton. Microbial ecology, 24(2):145–160, 1992.

[17] Ruud A Weusthuis, Jack T Pronk, PJ Van Den Broek, and JP Van Dijken. Chemostat cultivation as a tool for studies on sugar transport in yeasts. Microbiological reviews, 58(4):616–630, 1994.

[18] Thomas Pfeiffer, Stefan Schuster, and Sebastian Bonhoeffer. Cooperation and competition in the evolution of atp-producing pathways. Science, 292(5516):504–507, 2001.

[19] Robert E Beardmore, Ivana Gudelj, David A Lipson, and Laurence D Hurst. Metabolic trade-offs and the maintenance of the fittest and the flattest. Nature, 472(7343):342–346, 2011.

[20] Erik Postma, Cornelis Verduyn, W Alexander Scheffers, and Johannes P Van Dijken. Enzymic analysis of the crabtree effect in glucose-limited chemostat cultures of saccharomyces cerevisiae. Applied and environmental microbiology, 55(2):468–477, 1989.

[21] Annamaria Merico, Pavol Sulo, Jure Piškur, and Concetta Compagno. Fermentative lifestyle in yeasts belonging to the saccharomyces complex. The FEBS journal, 274(4):976–989, 2007.

[22] R Craig MacLean. The tragedy of the commons in microbial populations: insights from theoretical, comparative and experimental studies. Heredity, 100(5):471–477, 2008.

[23] Paul A Del Giorgio and Jonathan J Cole. Bacterial growth efficiency in natural aquatic systems. Annual Review of Ecology and Systematics, 29(1):503–541, 1998.

[24] Erik M Smith and Yves T Prairie. Bacterial metabolism and growth efficiency in lakes: the importance of phosphorus availability. Limnology and Oceanography, 49(1):137– 147, 2004.

[25] Craig A Carlson, Paul A Del Giorgio, and Gerhard J Herndl. Microbes and the dissipation of energy and respiration: from cells to ecosystems. Oceanography, 20(2):89–100, 2007.

[26] Vaibhhav Sinha and Seppe Kuehn. Hierarchical control of bacterial growth efficiency by substrate and taxonomy. *bioRxiv*, pages 2026–07, 2026.

[27] Dustin J Marshall, Hayley E Cameron, and Michel Loreau. Relationships between intrinsic population growth rate, carrying capacity and metabolism in microbial populations. The ISME Journal, 17(12):2140–2143, 2023.

[28] Simon P Hart, Robert P Freckleton, and Jonathan M Levine. How to quantify competitive ability. Journal of Ecology, 106(5):1902–1909, 2018.

[29] Eric R Pianka. r and k selection or b and d selection? The American Naturalist, 106(951):581–588, 1972.

[30] Meike T Wortel, Elad Noor, Michael Ferris, Frank J Bruggeman, and Wolfram Liebermeister. Metabolic enzyme cost explains variable trade-offs between microbial growth rate and yield. PLoS computational biology, 14(2):e1006010, 2018.

[31] Chuankai Cheng, Edward J O’brien, Douglas McCloskey, Jose Utrilla, Connor Olson, Ryan A LaCroix, Troy E Sandberg, Adam M Feist, Bernhard O Palsson, and Zachary A King. Laboratory evolution reveals a two-dimensional rate-yield tradeoff in microbial metabolism. PLoS computational biology, 15(6):e1007066, 2019.

[32] Steven A Frank. The trade-off between rate and yield in the design of microbial metabolism. Journal of evolutionary biology, 23(3):609–613, 2010.

[33] SJ Pirt. The maintenance energy of bacteria in growing cultures. Proceedings of the Royal Society of London. Series B. Biological Sciences, 163(991):224–231, 1965.

[34] HH Beeftink, RTJM Van der Heijden, and JJ Heijnen. Maintenance requirements: energy supply from simultaneous endogenous respiration and substrate consumption. FEMS Microbiology Ecology, 6(3):203–209, 1990.

[35] Gangsheng Wang and Wilfred M Post. A theoretical reassessment of microbial maintenance and implications for microbial ecology modeling. FEMS Microbiology Ecology, 81(3):610–617, 2012.

[36] Jana S Huisman, Martina Dal Bello, and Jeff Gore. Predictable shifts in microbial species composition lead to community-wide robustness to environmental stress. Nature Microbiology, pages 1–11, 2026.

[37] Sivan Pearl Mizrahi, Hyunseok Lee, Akshit Goyal, Erik Owen, and Jeff Gore. Structured interactions explain the absence of keystone species in synthetic microcosms. The ISME Journal, 19(1):wraf211, 2025.

[38] Matti Gralka, Shaul Pollak, and Otto X Cordero. Genome content predicts the carbon catabolic preferences of heterotrophic bacteria. Nature Microbiology, 8(10):1799– 1808, 2023.

[39] James B Russell and Gregory M Cook. Energetics of bacterial growth: balance of anabolic and catabolic reactions. Microbiological reviews, 59(1):48–62, 1995.

[40] Peter Van Bodegom. Microbial maintenance: a critical review on its quantification. Microbial ecology, 53(4):513–523, 2007.

[41] STEVEN S Dills, APRIL Apperson, MARY R Schmidt, and MILTON H Saier Jr. Carbohydrate transport in bacteria. Microbiological reviews, 44(3):385–418, 1980.

[42] KAZUMI Tanaka, SUMIKO Niiya, and TOMOFUSA Tsuchiya. Melibiose transport of escherichia coli. Journal of Bacteriology, 141(3):1031–1036, 1980.

[43] Christine Houssin, Nathalie Eynard, Emanuel Shechter, and Alexandre Ghazi. Effect of osmotic pressure on membrane energy-linked functions in escherichia coli. Biochimica et Biophysica Acta (BBA)-Bioenergetics, 1056(1):76–84, 1991.

[44] William G Roth, Mary P Leckie, and David N Dietzler. Osmotic stress drastically inhibits active transport of carbohydrates by escherichiacoli. Biochemical and biophysical research communications, 126(1):434–441, 1985.

[45] Guy Bunin. Ecological communities with lotka-volterra dynamics. Physical Review E, 95(4):042414, 2017.

[46] Matthieu Barbier, Claire De Mazancourt, Michel Loreau, and Guy Bunin. Fingerprints of high-dimensional coexistence in complex ecosystems. Physical Review X, 11(1):011009, 2021.

[47] Yuping Li, Dmitri A Petrov, and Gavin Sherlock. Single nucleotide mapping of trait space reveals pareto fronts that constrain adaptation. Nature ecology & evolution, 3(11):1539–1551, 2019.

[48] Maja Novak, Thomas Pfeiffer, Richard E Lenski, Uwe Sauer, and Sebastian Bonhoeffer. Experimental tests for an evolutionary trade-off between growth rate and yield in e. coli. The American Naturalist, 168(2):242–251, 2006.

[49] Olivia Kosterlitz, Elizabeth S Duan, Maya Abhyankar, Benjamin Kerr, and Eva M Top. Horizontal gene transfer by plasmids relaxes the evolutionary constraints of its own tradeoff. *bioRxiv*, pages 2026–05, 2026.

[50] Zehui Lao and Bei-Wen Ying. Growth dynamics of 3,909 escherichia coli single-gene knockouts in rich and minimal media. Scientific Data, 2026.

[51] Tomoya Baba, Takeshi Ara, Miki Hasegawa, Yuki Takai, Yoshiko Okumura, Miki Baba, Kirill A Datsenko, Masaru Tomita, Barry L Wanner, and Hirotada Mori. Construction of escherichia coli k-12 in-frame, single-gene knockout mutants: the keio collection. Molecular systems biology, 2(1):MSB4100050, 2006.

[52] Agata Jakubowska and Ryszard Korona. Epistasis for growth rate and total metabolic flux in yeast. PLoS One, 7(3):e33132, 2012.

[53] Xinzhu Wei and Jianzhi Zhang. Environment-dependent pleiotropic effects of mutations on the maximum growth rate r and carrying capacity k of population growth. PLoS Biology, 17(1):e3000121, 2019.

[54] Giulia Ghedini and Dustin J Marshall. Metabolic evolution in response to interspecific competition in a eukaryote. Current Biology, 33(14):2952–2961, 2023.

[55] Nora Underwood. Variation in and correlation between intrinsic rate of increase and carrying capacity. The American Naturalist, 169(1):136–141, 2007.

[56] Leo S Luckinbill. Selection and the r/k continuum in experimental populations of protozoa. The American Naturalist, 113(3):427–437, 1979.

[57] Paul B Rainey and Michael Travisano. Adaptive radiation in a heterogeneous environment. Nature, 394(6688):69–72, 1998.

[58] Richard Baran, Eoin L Brodie, Jazmine Mayberry-Lewis, Eric Hummel, Ulisses Nunes Da Rocha, Romy Chakraborty, Benjamin P Bowen, Ulas Karaoz, Hinsby Cadillo- Quiroz, Ferran Garcia-Pichel, et al. Exometabolite niche partitioning among sympatric soil bacteria. Nature communications, 6(1):8289, 2015.

[59] Jacob I Levine, Jonathan M Levine, Theo Gibbs, and Stephen W Pacala. Competition for water and species coexistence in phenologically structured annual plant communities. Ecology Letters, 25(5):1110–1125, 2022.

[60] Jacob I Levine, Stephen W Pacala, and Jonathan M Levine. Competition for time: Evidence for an overlooked, diversity-maintaining competitive mechanism. Ecology Letters, 27(3):e14422, 2024.

[61] Jennifer S Thaler, Scott H McArt, and Ian Kaplan. Compensatory mechanisms for ameliorating the fundamental trade-off between predator avoidance and foraging. Proceedings of the National Academy of Sciences, 109(30):12075–12080, 2012.

[62] Megan G Behringer, Wei-Chin Ho, Samuel F Miller, Sarah B Worthan, Zeer Cen, Ryan Stikeleather, and Michael Lynch. Trade-offs, trade-ups, and high mutational parallelism underlie microbial adaptation during extreme cycles of feast and famine. Current Biology, 34(7):1403–1413, 2024.

[63] Carlos Reding-Roman, Mark Hewlett, Sarah Duxbury, Fabio Gori, Ivana Gudelj, and Robert Beardmore. The unconstrained evolution of fast and efficient antibiotic-resistant bacterial genomes. Nature ecology & evolution, 1(3):0050, 2017.

[64] Christian Quast, Elmar Pruesse, Pelin Yilmaz, Jan Gerken, Timmy Schweer, Pablo Yarza, Jörg Peplies, and Frank Oliver Glöckner. The silva ribosomal rna gene database project: improved data processing and web-based tools. Nucleic acids research, 41(D1):D590–D596, 2012.

[65] Benjamin J Callahan, Paul J McMurdie, Michael J Rosen, Andrew W Han, Amy Jo A Johnson, and Susan P Holmes. Dada2: High-resolution sample inference from illumina amplicon data. Nature methods, 13(7):581–583, 2016.

[66] Stephen F Altschul, Warren Gish, Webb Miller, Eugene W Myers, and David J Lipman. Basic local alignment search tool. Journal of molecular biology, 215(3):403–410, 1990.

